# Anatomical, Physiological, and Functional Heterogeneity of the Dorsal Raphe Serotonin System

**DOI:** 10.1101/257378

**Authors:** Jing Ren, Drew Friedmann, Jing Xiong, Cindy D. Liu, Katherine E. DeLoach, Chen Ran, Albert Pu, Yanwen Sun, Brandon Weissbourd, Rachael L. Neve, Mark Horowitz, Liqun Luo

## Abstract

The dorsal raphe (DR) constitutes a major serotonergic input to the forebrain, and modulates diverse functions and brain states including mood, anxiety, and sensory and motor functions. Most functional studies to date have treated DR serotonin neurons as a single, homogeneous population. Using viral-genetic methods, we found that subcortical-vs. cortical-projecting serotonin neurons have distinct cell body distributions within the DR and different degrees of coexpressing a vesicular glutamate transporter. Further, the amygdala-and frontal cortex-projecting DR serotonin neurons have largely complementary whole-brain collateralization patterns, receive biased inputs from presynaptic partners, and exhibit opposite responses to aversive stimuli. Gain-and loss-of-function experiments suggest that amygdala-projecting DR serotonin neurons promote anxiety-like behavior, whereas frontal cortex-projecting neurons promote active coping in face of challenge. These results provide compelling evidence that the DR serotonin system contains parallel sub-systems that differ in input and output connectivity, physiological response properties, and behavioral functions.

## Introduction

The serotonin system powerfully modulates physiology and behavior in health and disease (Muller and Jacobs, 2010). Serotonin’s role in regulating emotional behavior has drawn particular attention as the serotonin system is the most widely used pharmacological target for treating depression and anxiety (Bandelow et al., 2008; Belmaker and Agam, 2008; Ravindran and Stein, 2010), and depression has become the leading cause of disability worldwide, with rates still on the rise (World Health Organization, 2017). However, a physiological and circuitry-based theory of how the serotonin system is organized to carry out its diverse functions remains elusive (Dayan and Huys, 2015; Muller and Jacobs, 2010). This poor understanding may partly account for the fact that the majority of antidepressant and anxiolytic drugs developed to target the 17 different serotonin receptors ultimately failed to reach the market (Berton and Nestler, 2006; Griebel and Holmes, 2013).

Serotoninergic fibers originate from only a few discrete nuclei in the brainstem but appear to innervate every part of the mammalian brain (Steinbusch, 1981). The largest serotonergic nucleus is the dorsal raphe nucleus (DR), which contains about 35% of the estimated 26,000 total serotonin-producing neurons in the mouse brain and is the major source of serotonergic innervation of the forebrain (Ishimura et al., 1988). Despite a large body of literature on functional perturbation using classic and modern techniques (Muller and Jacobs, 2010), we still lack a consensus as to the primary functions of the DR serotonin system. For example, studies on the effects of acute activation of DR serotonin neurons have reported divergent findings, including reinforcement (Liu et al., 2014), promotion of waiting for delayed reward rather than reinforcement (Fonseca et al., 2015; Miyazaki et al., 2012), promotion of anxiety-like behaviors and suppression of locomotion (Teissier et al., 2015; Urban et al., 2016), and suppression of locomotion without effects on reinforcement or anxiety-like behaviors (Correia et al., 2017). While different behavioral assays and activation methods may contribute to these conflicting results, they may also stem from treatment of the DR serotonin system as a monolithic whole.

Accumulating evidence points to electrophysiological, neurochemical, and molecular heterogeneity within the DR serotonin system (Calizo et al., 2011; Cohen et al., 2015; Fernandez et al., 2016; Hennessy et al., 2017; Okaty et al., 2015). Previous retrograde labeling studies also suggest that DR subregions may preferentially project to different target regions (reviewed in Waselus et al., 2011). In addition, whole-brain mapping of monosynaptic inputs to DR suggested heterogeneity of DR serotonin neurons with respect to presynaptic inputs (Weissbourd et al., 2014). Recent studies have begun to target subpopulations of DR serotonin neurons by utilizing optogenetic activation of serotoninergic terminals at specific targets (Marcinkiewcz et al., 2016) or genetic intersection combined with chemogenetic perturbations (Niederkofler et al., 2016). However, given the scale and complexity of the DR serotonin system, comprehensive characterizations that integrate anatomy, physiology, and function is essential for understanding how the serotonin system is organized to modulate diverse physiological and behavioral functions in health and disease. Using a combination of viral-genetic approaches, here we provide compelling evidence that the DR serotonin system contains parallel sub-systems that differ in sources of synaptic input, projection targets, physiological response properties, and behavioral functions.

## Results

### DR Serotonin Neurons that Project to Specific Targets Have Stereotyped Locations

To determine whether the spatial distribution of DR serotonin neuron cell bodies correlates with their projection targets, we performed retrograde tracing in combination with serotonin marker staining at the DR. We sampled eight brain regions reported by previous anterograde tracing studies to be heavily innervated by DR projections (Oh et al., 2014; Vertes, 1991), including those specifically from serotonin neurons (Allen Brain Atlas, 2017; http://connectivity.brain-map.org): the paraventricular hypothalamic nucleus (PVH), central amygdala (CeA), lateral habenula (LHb), dorsal portion of the lateral geniculate nucleus (dLGN), olfactory bulb (OB), orbitofrontal cortex (OFC), piriform cortex (PIR), and entorhinal cortex (ENT) (n=4 for each region). We injected ***HSV-Cre*** (Neve et al., 2005), which transduces neurons via their axon terminals, unilaterally into these regions in ***Ai14*** tdTomato Cre reporter mice (Madisen et al., 2010) (Figure 1A). Co-immunostaining DR sections with the serotonergic neuronal marker tryptophan hydroxylase 2 (Tph2), a biosynthetic enzyme for serotonin, revealed that serotonin neurons projecting to specific output sites have correspondingly stereotyped cell body locations in the DR. Specifically, serotonin neurons projecting to subcortical areas (PVH, CeA, LHb, and dLGN) tended to localize more in the dorsal DR, especially the dorsal wings (Figure 1B, top row). By contrast, serotonin neurons that project to the OB and three cortical areas (OFC, PIR, and ENT) preferentially localized in the ventral DR and were rarely found in the dorsal wing (Figure 1B, bottom row).

**Figure 1.**
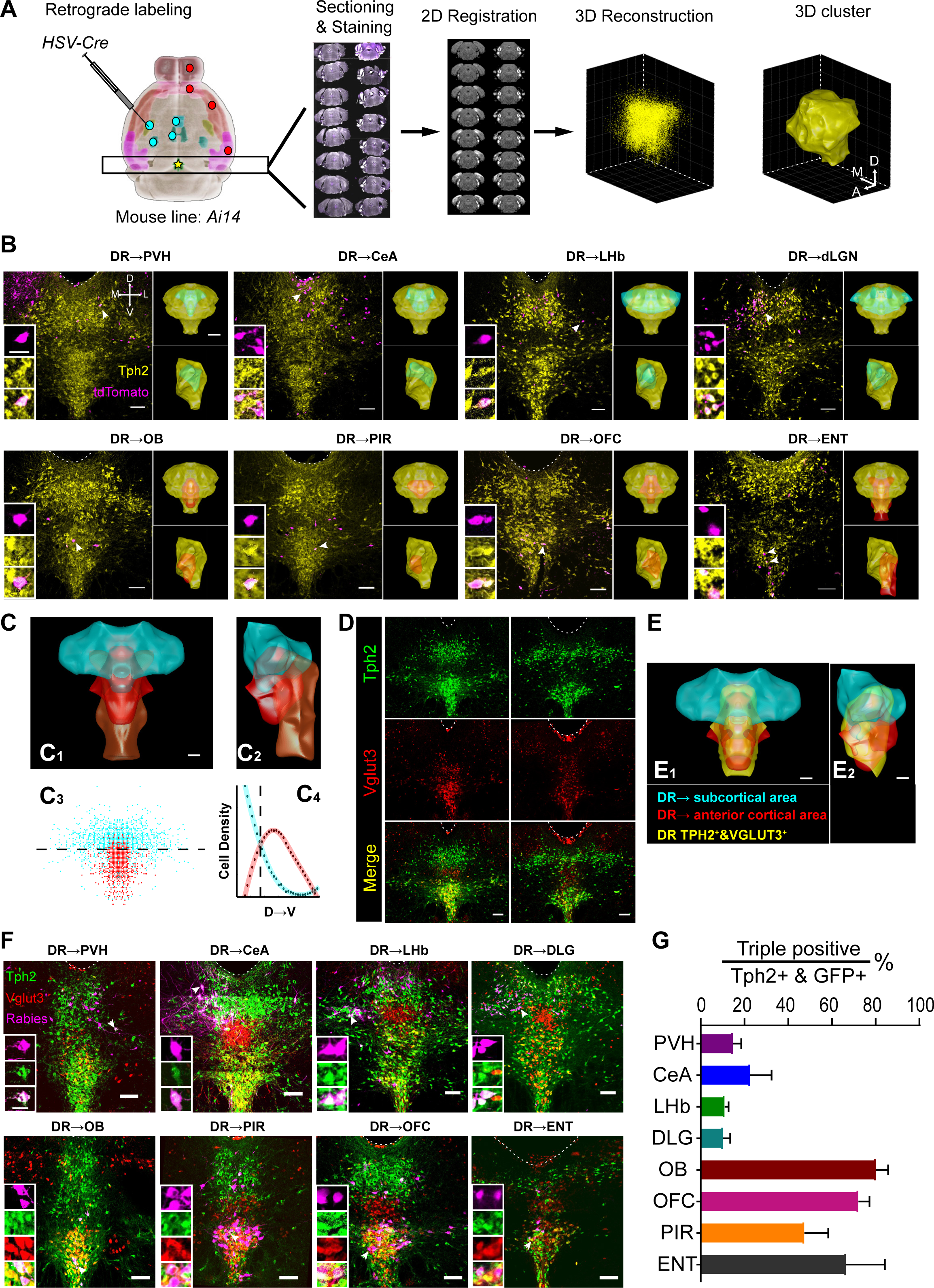
Spatial Organization of DR Serotonin Neurons with Respect to Axonal Projections and Vglut3 Co-expression. (A) Schematic of retrograde labeling and 3D reconstruction of DR serotonin neurons. ***HSV-Cre*** was injected into one of the eight brain regions (cyan and red dots represent injection sites) of ***Ai14*** mice: olfactory bulb (OB), orbitofrontal cortex (OFC), piriform cortex (PIR), entorhinal cortex (ENT), paraventricular hypothalamus (PVH), central amygdala (CeA), lateral habenula (LHb), and dorsal lateral geniculate nucleus (dLGN). Coronal sections were collected and kept in order. Anti-Tph2 staining was performed on sections containing DR (star). The positions of Tph2^+^/tdTomato^+^ cells were recorded in confocal images. Sections were each registered to a reference brain (Allen Institute for Brain Science, 2015) and reconstructed in 3D, thus assigning all Tph2^+^ cells (4698 ± 376.6 per brain) with Allen reference coordinates. DBSCAN was then applied for spatial clustering to generate a 3D surface of the DR serotonin system based on the location of Tph2^+^ neurons (STAR Methods). (B) Representative coronal confocal sections of the DR showing cells projecting to eight brain regions. Magenta shows the retrogradely labeled cells, yellow shows anti-Tph2 staining. Scale, 100 μm. Left insets, high-magnification images show neurons marked by arrows in the magenta, yellow, and both channels. Scale, 25 μm. Right insets (top, coronal view; bottom, sagittal view): yellow, cyan, and magenta structures represent 3D surface of the clusters of, respectively, all DR Tph2+ neurons; those that project to PVH, CeA, LHb, or dLGN; or those that project to OB, PIR, OFC or ENT. Scale, 500 μm. Dashed line, aqueduct (AQ). Cell numbers per brain, DR^Tph2^→PVH, 86 ± 9.4; DR^Tph2^→CeA, 115 ± 22.8; DR^Tph2^→LHb, 115 ± 9.9; DR^Tph2^→dLGN, 178 ± 66.0; DR^Tph2^→OB, 68 ± 8.0; DR^Tph2^→OFC, 127 ± 34.5; DR^Tph2^→PIR, 53 ± 15.4; DR^Tph2^→ENT, 72 ± 21.2. (C) Merged surface view of the DR^Tph2^→SC cluster (cyan), DR^Tph2^→AC cluster (red), and DR^Tph2^→ENT cluster (brown) in coronal (C_1_) and sagittal (C_2_) view. Scale, 200 μm. C_3_, coronal projection showing the location of individual cells from the DR^Tph2^→SC (cyan) and DR^Tph2^→AC (red) groups. Dashed line represent a plane 3742 μm ventral to the brain surface. C_4_, densities of DR^Tph2^→SC and DR^Tph2^→AC neurons along the dorsal-ventral axis. Dashed line shows where the two clusters share the same line density at 3742 μm ventral to the brain surface. (D) Representative coronal confocal sections of DR showing anti-Tph2 staining in ***Vglut3-Cre/Ai14*** mice (green), which express the Cre reporter tdTomato in Vglut3+ cells (red). (E) Coronal (E_1_) and sagittal (E_2_) view integrating projection-defined clusters (DR^Tph2^→SC, cyan; DR^Tph2^→AC, red) and the cluster of Tph2+&Vlgut3+ neurons (yellow, 1730 ± 219.8 neurons; n=3). Scale, 200 μm. (F) Representative coronal confocal sections of the DR showing retrogradely labeled neurons from eight brain regions (magenta, pseudo-colored from rabies-derived GFP), anti-Tph2 staining (green), and tdTomato from Vglut3-Cre+ neurons (red). Scale, 100 μm. Insets: magnified images showing the neurons indicated with arrows in individual channels. Scale, 25 μm. (G) Quantification of the proportion of GFP, Tph2, and Vglut3 triple positive neurons in GFP+/Tph2+ neurons for 8 projection brain regions listed on y-axis. (H) Axis label in this and all subsequent figures: A, anterior; P, posterior, D, dorsal; V, ventral; M, medial; L, lateral.

To systematically and quantitatively visualize the organization of these output-defined serotonin neurons, we developed an image registration algorithm that allowed us to register all DR-containing histological sections to the Allen Institute reference brain (Figure 1A, right panels; STAR Methods; Xiong et al., 2018). We then created clusters using the combined data from four brains with the same injection sites (Figure 1B, right insets). We found that serotonin neurons projecting to the four subcortical sites appeared largely overlapping in the dorsal DR. We combined these to produce a subcortical cluster (DR^Tph2^→SC) (Figure 1C). Likewise, serotonin neurons projecting to the OB, PIR, and OFC also exhibited considerable overlap, and we combined these to produce an anterior cortical cluster (DR^Tph2^→AC). ENT-projecting serotonin neurons tended to distribute more in caudal DR than other populations. DR^Tph2^→SC and DR^Tph2^→AC clusters preferentially occupied the dorsal and ventral DR, respectively, albeit with partial overlaps that accounted for 17.4% of DR^Tph2^→SC volume and 39.6% of DR^Tph2^→AC volume. This preferential distribution and partial overlap could also be seen in the cell body distribution along the dorsal-ventral (D-V) axis; 74% of DR^Tph2^→SC neurons were dorsal to the horizontal plane 3742 μm below the brain’s surface whereas 80% of DR^Tph2^→AC neurons were ventral to that plane (Figure 1C_3_-C_4_, Figure S1D-G). Together, these data demonstrate that cell bodies of DR serotonin neurons are organized according to their projection patterns (see Movie S1 for a summary).

### DR Serotonin Neurons that Co-express Vglut3 Preferentially Project to Cortical Regions

Previous findings suggest that ~60% of DR serotonin neurons co-express the vesicular glutamate transporter Vglut3 in the rat (Gras et al., 2002), and recent physiology studies suggest that these neurons co-release glutamate in the mouse (Liu et al., 2014; Sengupta et al., 2017). To investigate the distribution of dual-transmitter-containing neurons, we crossed ***Vglut3-Cre*** mice (Grimes et al., 2011) with the ***Ai14*** reporter and stained the DR sections with Tph2. We found that Vglut3 also subdivided the DR serotonin neurons roughly into dorsal and ventral compartments (Figure 1D). To visualize the 3D relationships between dual-transmitter-containing serotonin neurons (the DR^Vglut3&Tph2^ cluster) and our projection-defined serotonin neuron clusters, we performed double-labeling in three brains and registered them to the same reference atlas. We found that the DR^Vglut3&Tph2^ cluster largely matched with the spatial location of the DR^Tph2^→AC cluster rather than the DR^Tph2^→SC cluster (Movie S1). The combined DR^Vglut3&Tph2^ cluster had 66% ± 0.6% (mean ± SEM) overlap with the DR^Tph2^→AC but only 19.16% ± 1.1% overlap with the DR^Tph2^→SC cluster (Figure 1E).

These data suggest that projection and neurochemical patterns correlate, such that serotonin neurons that project to the OB and cortical regions more likely co-express Vglut3. To test this hypothesis, we injected GFP-expressing non-pseudotyped rabies virus as a retrograde tracer (Wickersham et al., 2007) into the eight aforementioned brain regions of ***Vglut3-Cre;Ai14*** animals, and determined the percentage of GFP^+^/tdTomato^+^/Tph2^+^ triple-labeled cells within the GFP^+^/Tph2^+^ double-labeled population (Figure 1F). Indeed, the Vglut3+ fractions in OB-and cortical-projecting serotonin neurons were high, whereas subcortical-projecting serotonin neurons were mostly Vglut3^−^ (Figure 1G). These data indicate that the DR serotonin system has a coordinated spatial and neurochemical organization with respect to its projection targets, although these subdivisions are not absolute.

### OFC- and CeA-projecting DR Serotonin Neurons Have Largely Complementary Collateralization Patterns

A key question regarding the anatomical organization of the serotonin system is to what extent serotonin neurons that project to one target region collateralize to other brain regions. Given the apparent distinction of cortical- and subcortical-projecting DR serotonin neurons (Figure 1), we chose to determine the complete collateralization patterns of OFC- and CeA- projecting ones as representative examples. We employed a recently developed intersectional strategy (Beier et al., 2015; Schwarz et al., 2015) to label DR serotonin neurons based on their projections. We utilized ***Sert-Cre*** mice (Gong et al., 2007), which express Cre recombinase in cells expressing the serotonin transporter (Sert), a serotonin neuron marker (Weissbourd et al., 2014). We injected axon-terminal-transducing AAV (Tervo et al., 2016) expressing Cre-dependent Flp recombinase ***(AAV_retro_-FLEx^loxP^-Flp)*** into the OFC or CeA, and AAVs expressing Flp-dependent membrane tethered GFP ***(AAV-FLEx^FLP^-mGFP)*** into the DR, of ***Sert-Cre*** mice (Figure 2A). We then employed iDISCO+ (Renier et al., 2016) to optically clear the brain in a whole-mount preparation, imaged the brains using a light-sheet microscope, and registered the imaged volume to the Allen Institute’s reference brain (Kim et al., 2015) to examine resultant axonal arborization patterns (STAR Methods).

**Figure 2.**
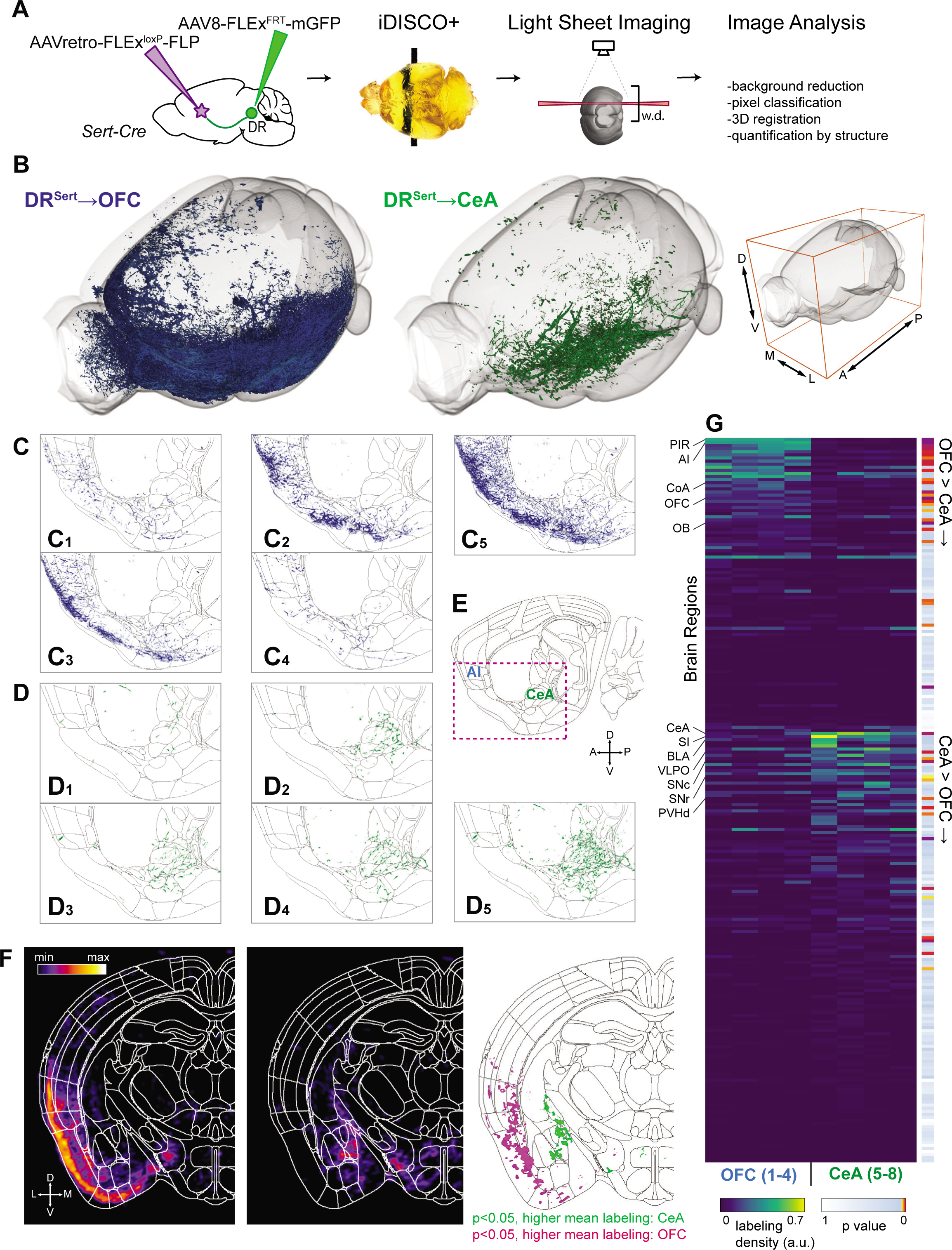
Distinct Collateralization Patterns of OFC- and CeA-Projecting DR Serotonin Neurons. (A) Schematic of viral-genetic tracing strategy and iDISCO+-based whole-brain 3D imaging. w.d., working distance of the light-sheet microscope objective. (B) Overview of axonal projections from one representative brain each from the DR^Sert^→OFC (blue) and DR^Sert^→CeA (green) groups. Whole-mount imaging included the entire left hemisphere and the medialmost ~650 μm of the right hemisphere. Viewing angle can be seen by the image on the right with axis labels. See Movie S2 for a 3D rendering. (C, D) Sagittal view of single 5-μm optical sections from eight individual brains registered to the Allen Institute common coordinate framework. Axons from DR^Sert^→OFC (C_1_-C_4;_ merged in C_5_) and DR^Sert^→CeA (D_1_-D_4_; merged in D_5_) neurons are shown in green and blue respectively. (E) Sagittal brain atlas image (10 μm) from the Allen Institute that encompasses CeA and anterior insular cortex (AI). The red box indicates the displayed region of (C) and (D). (F) Coronal density maps of DR^Sert^→CeA (left panel) and DR^Sert^→OFC (middle panel) projections generated by voxel-wise dilation of axons. Right panel shows a p-value map highlighting individual voxels with p < 0.05 between groups. See Movie S3 for the fly-through of these maps through the coronal series of the entire brain rostral to DR. (G) Heat map of relative labeling density (normalized to region volume and gross label content per brain) across 234 regions defined by the Allen Atlas. Regions are divided based on comparing mean intensity between groups, and values are then sorted by their second principal component. See Table S1 for the list of brain regions in the same sequence. AI, anterior insular cortex; CoA, cortical amygdala; SI, substantia innominata; BLA, basolateral amygdala; VLPO, ventrolateral preoptic nucleus; SNc, substantia nigra compacta; SNr, substantia nigra pars reticulata; PVHd, paraventricular hypothalamic nucleus, descending division.

We analyzed the whole-brain projection pattern of DR serotonin neurons labeled retrogradely from either the OFC or CeA (abbreviated as DR^Sert^→OFC and DR^Sert^→CeA neurons hereafter) in four brains each (Figure 2; Figure S2; Table S1). 3D rendering of the whole-brain projections from these two groups suggested that DR^Sert^→OFC and DR^Sert^→CeA axons exhibited complementary projection patterns, preferentially innervating superficial (cortical) and deep (subcortical) brain regions (Figure 2B; Movie S2). Projecting in a caudal-to-rostral direction and after passing through the lateral hypothalamus, the DR^Sert^→OFC axons appeared to split out a medial branch that travels dorsally to medial frontal cortex; the rest fan out laterally to the cortex, with some projecting posteriorly to the ENT, and others innervating the PIR and insular cortex anteriorly, eventually combining with the medial branch to jointly innervate other prefrontal areas and the OB (Movies S2). By contrast, the DR^Sert^→CeA axons intensely innervated most amygdala nuclei and several hypothalamic nuclei, notably the ventrolateral preoptic nucleus and PVH. These axons also innervated the ventral part of the bed nucleus of stria terminalis (BNST), and terminated in the anterior substantia innominata (Movie S2; Table S1). Consistent with our retrograde tracing experiments in Figure 1, DR^Sert^→OFC axons did not innervate most forebrain subcortical regions, whereas DR^Sert^→CeA axons rarely invaded any cortical areas.

Close examination of axon patterns in thin optical sections supported the notion that the projection patterns of DR^Sert^→OFC and DR^Sert^→CeA neurons were largely complementary, and also revealed the stereotypy of individual brains from the same group despite large differences in labeling intensity (Figure 2C-E). As additional examples, the OFC and OB were intensively innervated by DR^Sert^→OFC axons but devoid of DR^Sert^→CeA axons (Figure S2A). By contrast, the CeA, PVH, BNST, and substantia nigra were innervated by DR^Sert^→CeA axons but devoid of DR^Sert^→OFC axons (Figure 2G; Figure S2B). DR^Sert^→CeA axons also innervated the lateral amygdala, basolateral amygdala, and intercalated nucleus of the amygdala (Table S1). However, the nearby cortical amygdala was innervated mostly by DR^Sert^→OFC axons rather than DR^Sert^ →CeA axons (Figure S2A).

Whole-brain quantitative and statistical analyses of DR^Sert^→OFC and DR^Sert^→CeA axonal projections validated that the innervation patterns were largely complementary throughout the brain (Figure 2F-G; Movie S3; Table S1). These analyses also indicated that many known targets of DR serotonin neurons (Allen Brain Atlas, 2017; Azmitia and Segal, 1978; Vertes, 1991) were not innervated by either of these subpopulations. These include most of the striatum, thalamus, hypothalamus, somatosensory and motor cortex, and dorsal parts of the BNST (Table S1). Thus, the DR serotonin system contains at least two, and likely more, parallel sub-systems with distinct innervation patterns.

### OFC- and CeA-projecting DR serotonin neurons receive biased input from specific nuclei

We and others have previously shown that DR serotonin neurons as a whole receive monosynaptic inputs from diverse brain regions (Ogawa et al., 2014; Pollak Dorocic et al., 2014; Weissbourd et al., 2014). Given their parallel output organization, we next used the viral-genetic strategy cTRIO (cell-type-specific tracing the relationship between input and output) (Schwarz et al., 2015) to investigate whether DR^Sert^→OFC and DR^Sert^→CeA neurons receive input from similar or different brain regions. We injected ***HSV-STOP^fox^-Flp*** unilaterally into the OFC or CeA, and AAVs carrying Flp-dependent constructs expressing TVA-mCherry fusion protein and rabies glycoprotein into the DR of ***Sert-Cre*** mice. We then injected EnvA-pseudotyped, glycoprotein-deleted, GFP-expressing rabies virus (RWG) into the DR. Thus, only Sert-Cre^+^ neurons that project to the OFC or CeA could become starter cells for RWG-mediated transsynaptic tracing (Figure 3A).

**Figure 3.**
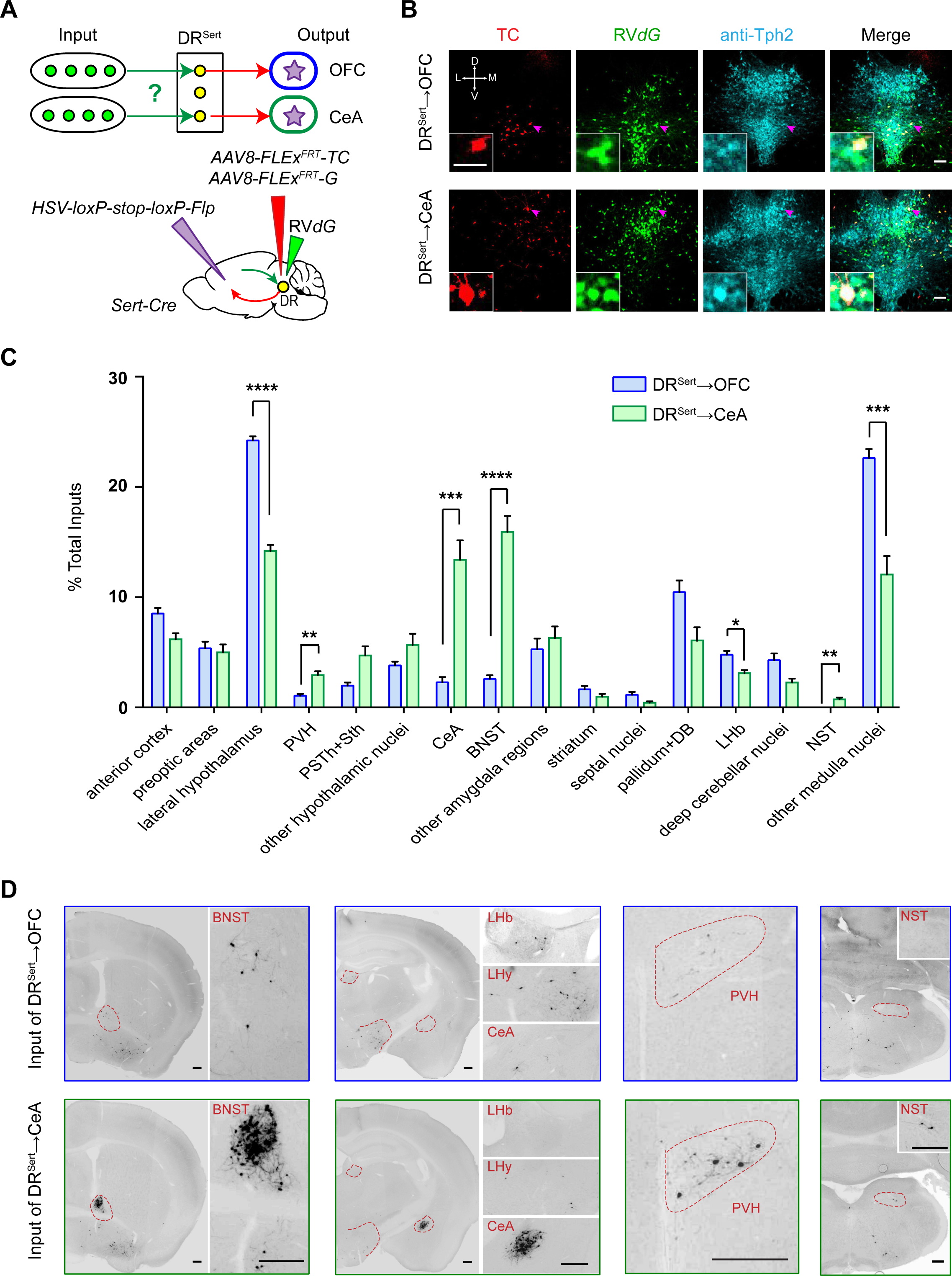
cTRIO Analysis Reveals Biased Input Distributions for OFC- and CeA-projecting DR Serotonin Neurons. (A) Schematic of cTRIO experiments applied to the DR. ***HSV-STOP^flox^-Flp*** was injected into either OFC or CeA of ***Sert-Cre*** mice, and AAVs expressing Flp-dependent TVA-mCherry (TC) and rabies glycoprotein (G) were injected into the DR. Two weeks later, RV***dG***-GFP was injected into the DR to initiate retrograde transsynaptic tracing. (B) Confocal images of coronal sections containing the DR, showing OFC- and CeA-projecting serotoninergic starter cells (starter cell numbers for DR^Sert^→OFC: 73 ± 8.3, n=9 mice; DR^Sert^→CeA: 96 ± 13.4, n=8 mice). Red shows TC expression, green shows GFP expression, and cyan shows anti-Tph2 staining. Scale, 100 μm. Insets: high magnification images of neurons pointed by arrows. Scale, 50 μm. (C) Quantification of whole-brain inputs to DR^Sert^→OFC and DR^Sert^→CeA neurons (n=9, 8). Y-axis presents percentage of total inputs of each brain, and x-axis lists brain regions. Error bars represent SEM *p<0.05, **p<0.01, ***p<0.001 and ****p<0.0001 (multiple t-tests with Holm-Sidak correction). PSTh, parasubthalamic nucleus; Sth, subthalamic nucleus; BNST, bed nucleus of the stria terminalis; DB, nucleus of the diagonal band; NST, nucleus of solitary tract. (D) Representative GFP labeled ipsilateral input cells to DR^Sert^→OFC and DR^Sert^→CeA neurons. LHy, lateral hypothalamus; DCN, deep cerebellum nuclei. Scale, 250 μm. (E) See Figure 1 for other abbreviations, and Figures S3 and Table S2 for related data.

Tph2 staining verified that starter cells were predominantly serotonin neurons (97% ± 2% and 94% ± 4% for DR^Sert^→OFC and DR^Sert^→CeA neurons, respectively; Figure 3B), and the location of starter cells was consistent with our previous findings (Figure 1). While thousands of long-range input cells were identified in each experimental group, few GFP-labeled cells were found in the two control groups for each experimental group: those without AAV expressing rabies glycoprotein and those using wild-type instead of ***Sert-Cre*** mice (Figure S3A-C). These data confirmed that long-range tracing of inputs was cell-type specific.

To determine the long-range presynaptic partners for each group of projection-defined DR serotonin neurons, we first divided each brain into 35 regions of interest (six cortical areas, 11 hypothalamic nuclei, four amygdala regions, four basal ganglia regions, ten medulla nuclei, thalamus, and cerebellum; Table S2) (Weissbourd et al., 2014). We counted the number of cells in each region from serial coronal sections, omitting regions around the DR due to possible local background labeling (Figure S3B). While presynaptic inputs to these two DR serotonin subpopulations originated from similar brain regions, there were striking quantitative differences. Specifically, DR^Sert^→OFC neurons received proportionally more input from lateral hypothalamus, LHb, and the majority of medulla nuclei. By contrast, DR^Sert^→CeA neurons received significantly more input from CeA itself, BNST, PVH, and nucleus of the solitary tract (Figure 3C-D; Figure S3D). The marked enrichment of CeA input to DR^Sert^→CeA neurons suggests strong DR-CeA reciprocal connectivity.

In summary, DR^Sert^→OFC and DR^Sert^→CeA neurons have largely complementary collateralization patterns (Figure 2) and receive quantitatively biased input from specific brain regions (Figure 3). In the rest of the study, we explore whether DR ^Sert^→OFC and DR^Sert^→CeA neurons also exhibit different physiological response properties and behavioral functions.

### OFC- and CeA-projecting DR Serotonin Neurons Are Both Activated by Reward But Show Opposite Responses to Punishment

Previous studies showed that when treated as a single group, DR serotonin neurons were activated during reward consumption in freely moving mice (Li et al., 2016). However, singleunit recordings in head-fixed mice revealed heterogeneous responses of DR serotonin neurons to reward and punishment (Cohen et al., 2015), although the origin of the heterogeneity is unclear. Since DR^Sert^→-OFC and DR^Sert^→CeA neurons receive biased presynaptic inputs from different brain regions (Figure 3), it is possible that they may respond differently to reward and punishment. To test this, we combined our viral-genetic strategy for accessing these different serotonin neuron subpopulations with fiber photometry (Gunaydin et al., 2014). We expressed a genetically encoded Ca^2+^ indicator GCaMP6m (Chen et al., 2013) in DR^Sert^→OFC or DR^Sert^→CeA neurons by bilaterally injecting ***AAV_retro_-FLEx-Flp*** into the projection site along with an AAV expressing Flp-dependent GCaMP6m into the DR of ***Sert-Cre*** mice. As a control for these projection-specific DR serotonin neurons, we also expressed Cre-dependent GCaMP6m in the DR of ***Sert-Cre*** mice (“DR^Sert^ group”). We implanted an optical fiber into the DR at the GCaMP6m injection site through which we delivered excitation and control light to monitor the activity of different serotonin neuron groups (Figure 4 A_1_-C_1_) (Allen et al., 2017). We verified GCaMP6m expression and recording sites via ***post hoc*** histology (Figure 4A_2_-C_2_; Figure S4A_1_-C_1_).

**Figure 4.**
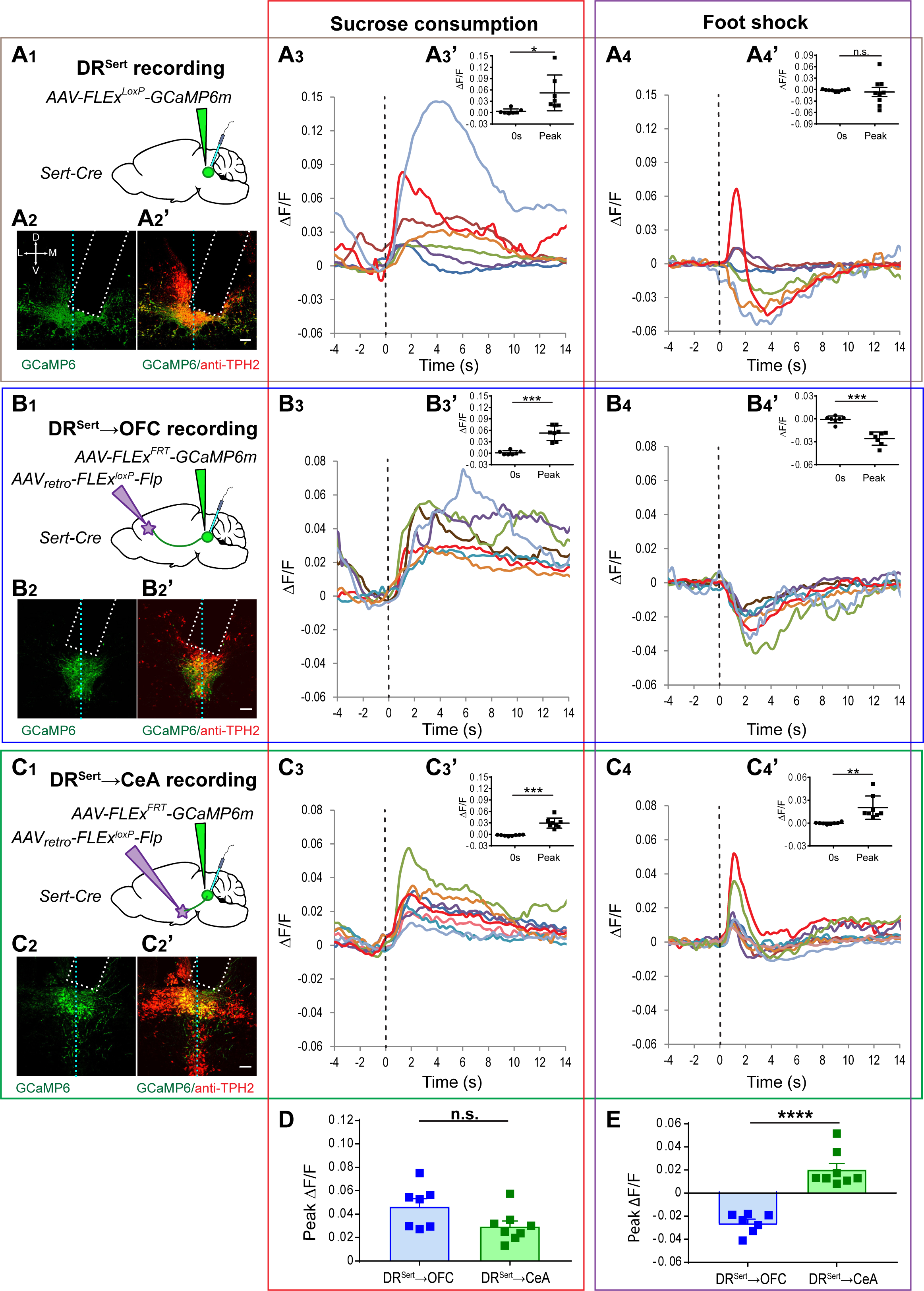
OFC- and CeA-projecting DR Serotonin Neurons Are Both Activated by Reward but Show Opposite Responses to Punishment. Fiber photometry recordings were performed on DR^Sert^ (A), DR^Sert^→OFC (B) and DR^Sert^→CeA neurons. (A_1_, B_1_, C_1_) Schematic of viral injection and optical fiber (blue) implantation. (A**2**, B**2**, C**2**) Confocal images of coronal sections showing fiber optic placement (dotted rectangle), and the expression of GCaMP6m (green) with Tph2 staining (red, A_2_’, B_2_’, C_2_’) in the DR. Vertical dashed lines represent the midline. Scale, 100 μm. (Estimate of GCaMP6m+ serotonin neurons under optical fiber: DR^Sert^ group, 204 ± 39.0, n=7 mice; DR^Sert^→OFC group, 112 ± 28.0, n=7 mice; DR^Sert^→CeA group, 115 ± 22.1, n=8 mice). (A_3_, B_3_, C_3_) Mean responses of individual mice to sucrose consumption. Time 0 is aligned to lick initiation (vertical dashed line). Red traces correspond to the mice shown in (A_2_, B_2_, C_2_) and Figure S4. (A_3_’, B_3_’, C_3_’) Group data showing quantification of the peak ***ΔF/F*** recorded during sucrose water licking comparing to the ***ΔF/F*** at time 0. (A_4_, B_4_, C_4_) Mean responses of individual mice from the three groups to electrical shock. Time 0 is aligned to onset of 1-sec electric shock delivery. (A_4_’, B_4_’, C_4_’) Group data showing quantification of the peak ***ΔF/F*** (negative or positive extreme) recorded after electric shock delivery comparing to the ***ΔF/F*** at time 0. (D) Quantification of the peak ***ΔF/F*** recorded during sucrose water licking from DR^Sert^→OFC and DR^Sert^→CeA neurons. (E) Quantification of the peak ***ΔF/F*** (negative or positive extreme) recorded after electric shock delivery comparing DR^Sert^→OFC and DR^Sert^→CeA neurons. Error bars represent s.e.m; n.s., not significant; *p<0.05; **p<0.01; ***p<0.001; ***p<0.001; ****p<0.0001 (n=7, 8 mice for DR^Sert^→OFC and DR^Sert^→CeA groups, respectively; two-tail paired t-test for A-C and unpaired t-test for D-E). See also Figure S4.

To record reward responses, we trained mice to lever press for a sucrose water reward in a fixed-ratio paradigm. Each lever press led to one unit of sucrose water delivered from a nearby port, and water-restricted mice were allowed free access during the recording. Consistent with a previous report (Li et al., 2016), all seven recordings from the DR^Sert^ group were activated by the onset of licking, and the evoked activity persisted during reward consumption period (Figure 4A_3_, A_3_’). The DR^Sert^→OFC and DR^Sert^→CeA groups were similarly activated (Figure 4B_3_, B_3_’, C_3_, C_3_’, D).

Next, we recorded responses to punishment by subjecting the same set of animals to mild electrical shocks applied to their feet through an electrified floor. Remarkably, DR^Sert^→OFC and DR^Sert^→CeA neurons responded to foot-shock in opposite fashion. All seven mice from the DR^Sert^→OFC group showed a long-lasting reduction of Ca^2+^ signals (~10 sec) during and after the 1-sec foot-shock (Figure 4B_4_, B_4_’, E). By contrast, in all eight mice from the DR^Sert^→CeA group, foot-shock induced a transient elevation (~2 sec) of Ca^2+^ signals, followed by a small depression in a subset (Figure C_4_, C_4_’, E). The seven mice from the DR^Sert^ group showed more varied responses, including one that exhibited a clear biphasic response composed of transient elevation followed by long-lasting depression (Figure 4A_4_, A_4_’).

To test the possibility that the foot-shock responses were related to novelty of the stimulus, we also recorded responses to unpredicted tones delivered the day prior to the footshock test. We did not observe any tone-induced responses (data not shown), suggesting that the foot-shock responses were caused by aversive stimuli rather than novelty. Altogether, our data indicate that DR^Sert^→OFC and DR^Sert^→CeA neurons respond similarly to reward but oppositely to punishment, and suggest that previous heterogeneous physiological responses of DR serotonin neurons to reward and punishment (Cohen et al., 2015) could also reflect recordings from projection-specific subpopulations.

### Both OFC- and CeA-projecting DR Serotonin Neurons suppress Locomotion

To investigate the behavioral functions of DR^Sert^→OFC and DR^Sert^→CeA neurons, we employed two complementary approaches. In the first (gain-of-function) approach, we expressed the hM3Dq chemogenetic activator (Armbruster et al., 2007) in DR^Sert^→OFC or DR^Sert^→CeA neurons by unilaterally injecting ***AAV_retro_-FLEx-Flp*** into the OFC or CeA of ***Sert-Cre*** mice, and Flp-dependent hM3Dq (Beier et al., 2017) into the DR in the experimental group. In the two control groups for each experimental group, we replaced either hM3Dq with GFP or ***Sert-Cre*** mice with wild-type mice (Figure 5A_1_, B_1_). ***Post hoc*** histology confirmed that hM3Dq expression was Cre-dependent and cells’ locations were consistent with previous results (Figure 5A_2_, B_2_). In the second (loss-of-function) approach, we conditionally knocked-out the ***Tph2*** gene from DR^Sert^→OFC or DR^Sert^→CeA neurons by bilaterally injecting ***AAV_retro_-Cre-2A-GFP*** into the projection site of ***Tph2^flox/flox^*** mice (Wu et al., 2012) 17 days prior to the onset of behavioral tests. In the control group, we injected ***AAV_retro_-GFP*** instead (Figure 5C_1_, D_1_). Immunostaining of ***Tph2^flox/flox^*** mice showed that GFP+ DR neurons lacked Tph2 protein (Figure 5C_2_, D_2_), confirming the effectiveness of the viral-mediated knockout.

**Figure 5.**
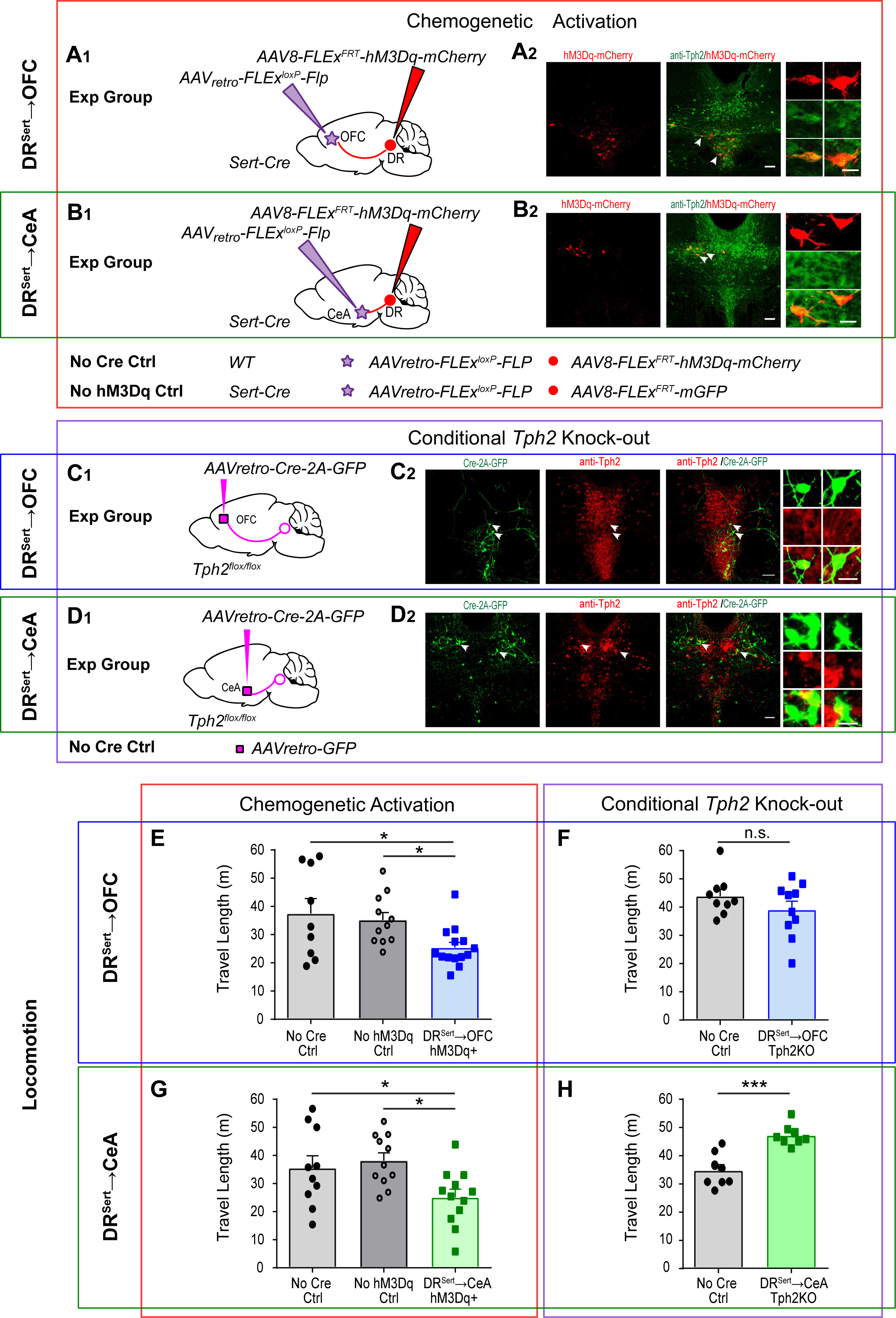
Chemogenetic Activation and Conditional ***Tph2*** Knockout Reveal that Both PFC- and CeA-projecting DR Serotonin Neurons suppress Locomotion. (A, B) Chemogenetic activation of DR^Sert^→OFC (A) and DR^Sert^→CeA (B) neurons. Exp, experimental. Ctrl, control. (A_1_, B_1_) Schematic for experimental groups. Conditions for the two controls are listed below. (A_2_, B_2_) Confocal images of coronal sections showing the expression of hM3Dq-2A-mCherry (red), all of which were double-labeled with Tph2 staining (green) in the DR. Dotted lines are borders of the aqueduct. Scale, 100 μm. Right panel: high magnified images showing neurons labeled by arrows. Scale, 50 μm. (Cell numbers, DR^Sert^→OFC group, 83 ± 4.8, n=14; DR^Sert^→CeA group, 93 ± 7.7, n=12) (C, D) DR^Sert^→OFC (C) and DR^Sert^→CeA (D) neurons were Tph2 depleted by injecting ***AAV_retro_-Cre*** into OFC (c_1_) or CeA (D_1_) of the conditional knockout ***Tph2^flox/fox^*** line bilaterally. (C**2**, D**2**) Confocal images of coronal sections showing the expression of Cre-2A-GFP (green), almost all of which were negative from Tph2 staining (red) in the DR. (Cells counted: Exp DR^Sert^→OFC group, 272 ± 67.9, 98.6% ± 0.43% were Tph2 negative, n=3 mice; Exp DR^Sert^→CeA group, 331 ± 76.6, 98.7% ± 0.33% were Tph2 negative, n=3 mice. Ctrl DR^Sert^→OFC group, 160 ± 18.5, 16.7% ± 0.18% were Tph2 negative, n=3 mice; Ctrl DR^Sert^→CeA group, 177 ± 42.2, 39.2% ± 0.16% were Tph2 negative). Dotted lines are borders of the aqueduct. Scale, 100 μm. Right panels: high magnification images showing neurons pointed by arrows. Scale, 25 μm. (E) Chemogenetic activation of hM3Dq+ DR^Sert^→OFC neurons decreases distance traveled (oneway ANOVA followed by multiple t-tests, Holm-Sidak correction; n= 9, 11, 14). *p<0.05. (F) Conditionally knocking out ***Tph2*** from DR^Sert^→OFC neurons does not have a significant effect on distance traveled (two-tail unpaired t-test; n=9, 10). (G) Activation of hM3Dq+ DR^Sert^→CeA neurons decreases distance traveled (one-way ANOVA followed by multiple t-test, Holm-Sidak correction; n= 10, 11, 12). *p<0.05. (H) Conditionally knocking out ***Tph2*** from DR^Sert^→CeA neurons increases distance traveled (two-tail unpaired t-test; n=8, 8). ***p<0.001. Error bars, SEM

Because this is, to our knowledge, the first functional investigation of projection-specific serotonin neuron subpopulations, we carried out a broad screen by subjecting the same sets of mice to a series of behavioral paradigms known to engage the DR serotonin system (STAR Methods) (Teissier et al., 2015). We first quantified locomotion in the open field and found that chemogenetic activation of both DR^Sert^→OFC and DR^Sert^→CeA neurons significantly decreased locomotion compared with controls (Figure 5E, G). Conversely, Tph2 depletion from DR^Sert^→CeA neurons significantly increased locomotion compared to controls, suggesting that serotonin is responsible for the locomotion suppression promoted by the DR→CeA neurons (Figure 5H). However, Tph2 depletion from DR^Sert^→OFC neurons did not affect locomotion (Figure 5F); one possible explanation is that the effect caused by activating this subpopulation involve other neurotransmitter(s) such as glutamate, as most DR^Sert^→OFC neurons were Vglut3+ (Figure 1G).

### CeA- but not OFC-projecting DR Serotonin Neurons Promote Anxiety-like Behavior

Next, we evaluated the effects of manipulating DR^Sert^→OFC and DR^Sert^→CeA neurons on anxiety-like behavior. Excessive avoidance of the center in the open field (Prut and Belzung, 2003) or the open arms of the elevated plus maze (EPM) (Walf and Frye, 2007) are widely used as indicators for anxiety-like behavior. We found that activation of DR^Sert^→OFC neurons did not affect center entry or center time in the open field, or open-arm entry or time in open arms of the EPM (Figure 6 A, E). However, Tph2 depletion from DR^Sert^→OFC neurons caused a significant decrease in center time and center entry compared with controls (Figure 6B), as well as a significant decrease in open-arm time in the EPM (Figure 6F). These data suggest that serotonin release in the DR→OFC projection has an anxiolytic effect (see Discussion).

**Figure 6.**
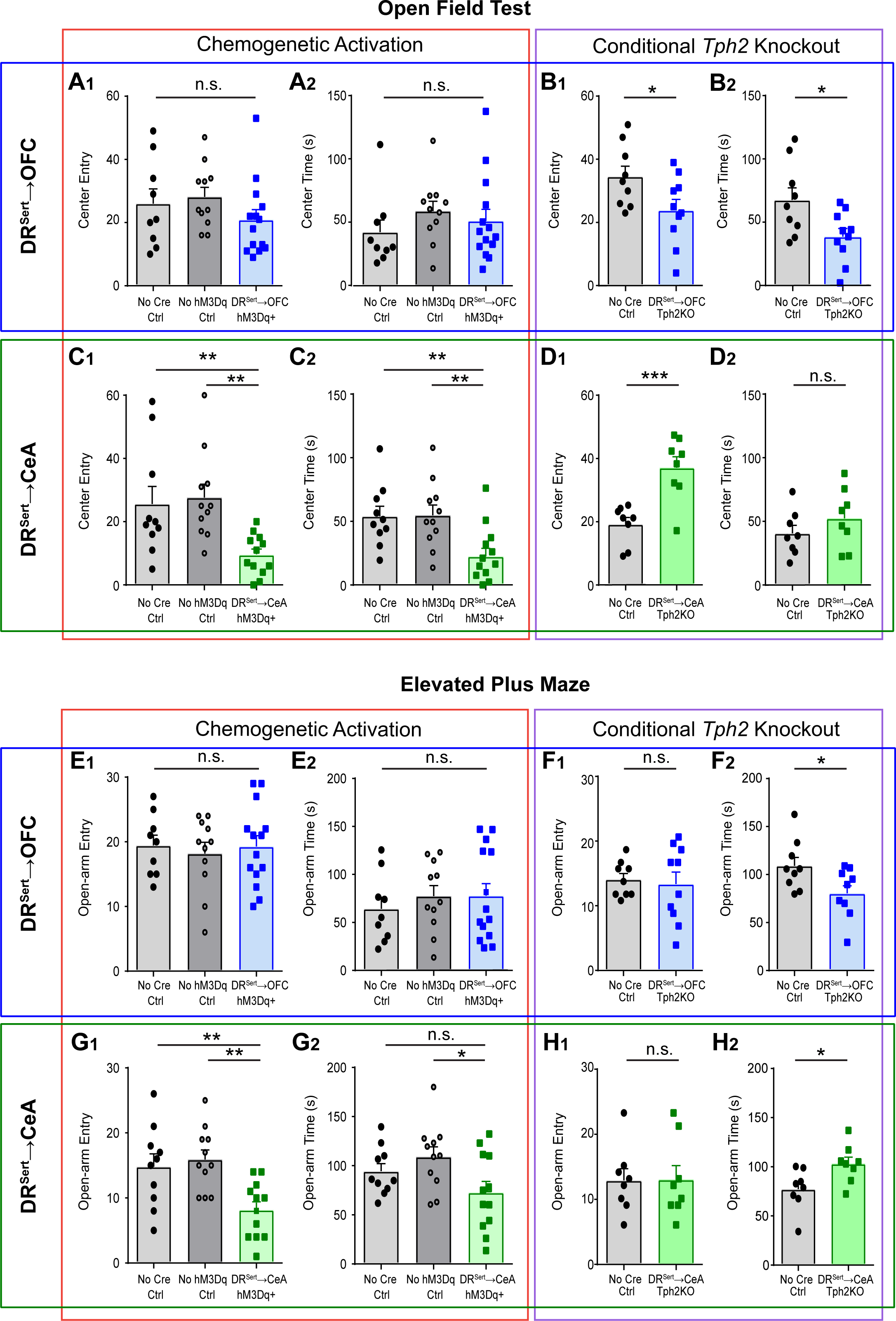
CeA- but Not PFC-projecting DR Serotonin Neurons Promote an Anxiety-like State. (A) Chemogenetic activation of hM3Dq+ DR^Sert^→OFC neurons does not affect the number of entries to the center (A_1_) or time spent in the center (A_2_) of the open field (one-way ANOVA; n= 9, 11, 14). (B) Conditionally knocking out ***Tph2*** from DR^Sert^→OFC neurons decreases the number of entries to the center of the open field (B_1_) and the time spent in the center (B_2_) (two-tail unpaired t-test; n=9, 10). (C) Activation of hM3Dq+ DR^Sert^→CeA neurons decreases the number of entries to the center of the open field (C_1_) and the time spent in the center (C_2_) (one-way ANOVA followed by multiple t-tests with Holm-Sidak correction; n= 10, 11, 12). (D) Conditionally knocking out ***Tph2*** from DR^Sert^→CeA neurons increases the number of entries to the center of the open field (D_1_), but not the time spent in the center (D_2_) (two-tail unpaired t-test; n=8, 8). (E) Activation of hM3Dq+ DR^Sert^→OFC neurons does not affect the number of entries to the open arm (E_1_) or the time spent in the open arm in the EPM (E_2_) (one-way ANOVA; n= 9, 11, 14). (F) Conditionally knocking out ***Tph2*** from DR^Sert^→OFC neurons does not affect the number of entries to the open arm (F_1_), but increases the time spent in the open arm in EPM (F_2_) (two-tail unpaired t-test; n=9, 10). (G) Activation of hM3Dq+ DR^Sert^→CeA neurons decreases the number of entries to the open arm (G_1_), and decreases the time spent in the open arm comparing to the no Cre Ctrl (G_2_) (one-way ANOVA followed by multiple t-tests, Holm-Sidak correction; n= 10, 11, 12). (H) Conditionally knocking out ***Tph2*** from DR^Sert^→CeA neurons does not affect the number of entries to the open arm (H_1_), but increased the time spent in the open arm (H_2_) (two-tail unpaired t-test; n=8, 8). *p<0.05; **p<0.01; ***p<0.001. Error bars, SEM

By contrast, activation of DR^Sert^→CeA neurons promoted anxiety-like behavior, resulting in significantly decreased center entry and center time in the open field compared to both control groups (Figure 6C). In EPM, activation of these neurons significantly decreased open arm entry compared to both control groups, and significantly decreased open arm time to one control group (Figure 6G). Accordingly, Tph2 depletion from DR^Sert^→CeA neurons appeared anxiolytic, causing a significant increase in center entry in the open field (Figure 6D) and open arm time in EPM (Figure 6H). Taken together, these data suggest that DR^Sert^→CeA neurons are anxiogenic, and that DR^Sert^→CeA and DR^Sert^→-OFC neurons have distinct functions in regulating anxiety-like behavior.

We also used fear conditioning to test the effect of manipulating DR^Sert^→CeA and DR^Sert^→-OFC neurons. We found that neither activation nor Tph2 depletion of DR^Sert^→OFC neurons affected fear learning and 1-day memory recall (Figure S5A, B, E, F). By contrast, activation of DR^Sert^→CeA neurons significantly increased overall freezing time to a conditioned tone in operant fear learning and 1-day memory retrieval session (Figure S5C, D). However, fear learning and memory were not affected by Tph2 depletion in DR^Sert^→CeA neurons (Figure S5G, H), suggesting that other pathways may compensate for the loss of DR^Sert^→CeA activity in fear learning and memory.

### OFC- but not CeA-projecting DR Serotonin Neurons Enhance Escape Behavior in Forced-Swim Test

Finally, we asked whether DR^Sert^→CeA and DR^Sert^→OFC neurons could modulate coping behavior a 2-day forced-swim test. Immobility and struggle (escape behavior) in the forced-swim test are often used as indicatives of passive and active coping in face of challenge, respectively (Petit-Demouliere et al., 2005). During chemogenetic experiments, CNO was administrated only before the day-2 test. While chemogenetic activation of DR^Sert^→CeA neurons did not affect immobility (Figure 7C), Tph2 depletion significantly enhanced escape behavior (reduced immobility) during the forced-swim test (Figure 7D), indicating that DR^Sert^→CeA neurons inhibit escape behavior, although we cannot rule out this being a secondary consequence of movement suppression (Figure 5H).

**Figure 7.**
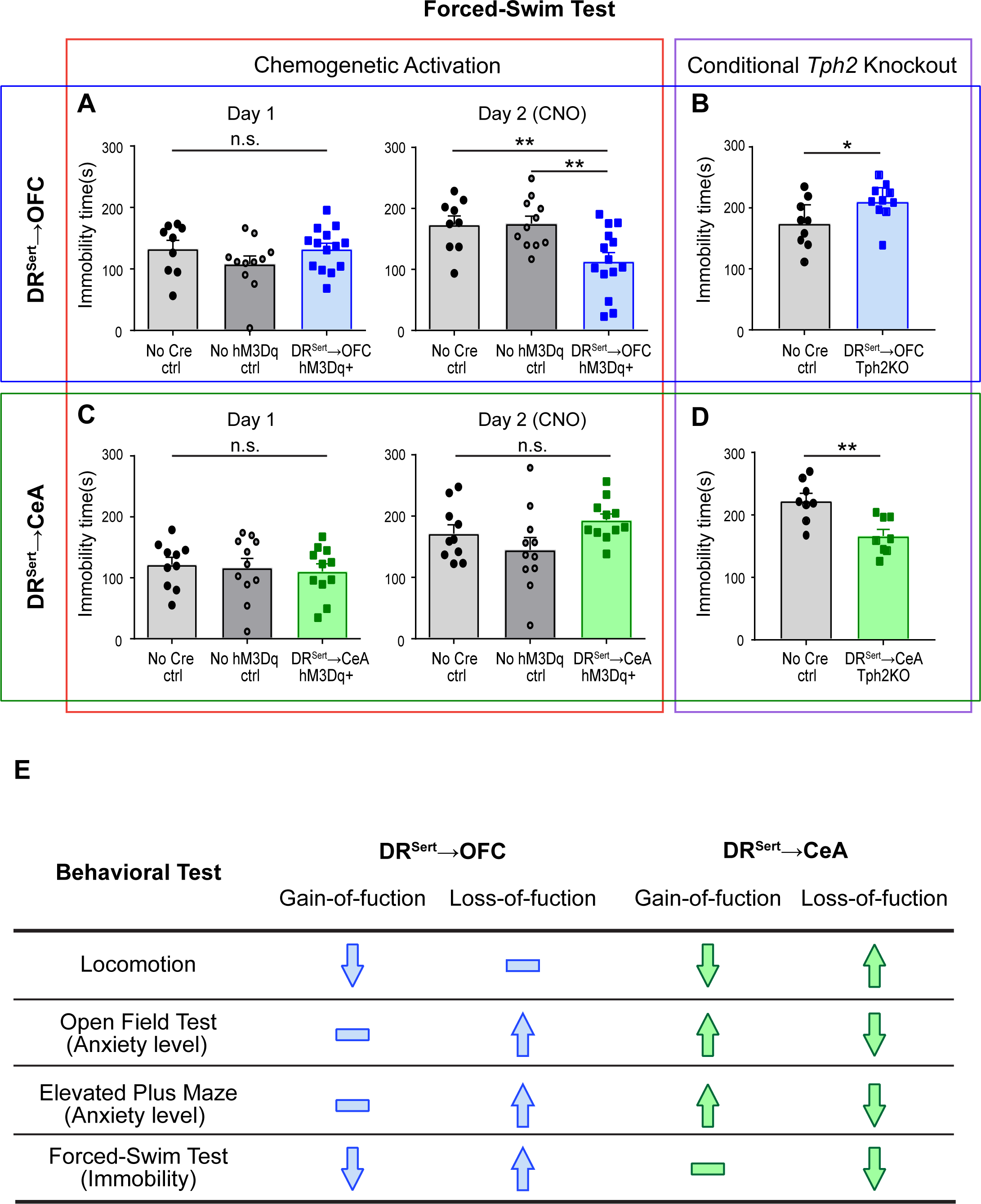
OFC-but Not CeA-projecting DR Serotonin Neurons Promote Escape Behavior in the Forced-Swim Test. (A) Activation of hM3Dq+ DR^Sert^→OFC neurons does not affect the immobility time in forced-swim test (FST) on Day 1 training session, but decreases the immobility time on day-2 FST testing session (one-way ANOVA followed by multiple t-tests, Holm-Sidak correction; n= 9, 11, 14). **p<0.01. (B) Conditionally knocking out ***Tph2*** from DR^Sert^→OFC neurons increases the immobility time in FST (two-tail unpaired t-test; n=9, 10). *p<0.05. (C) Activation of hM3Dq+ DR^Sert^→CeA neurons does not affect the immobility time in FST (one-way ANOVA; n= 10, 11, 11). (D) Conditionally knocking out ***Tph2*** from DR^Sert^→CeA neurons decreases the immobility time in FST (two-tail unpaired t-test; n=8, 8). **p<0.01. Error bars, SEM (E) Summary of gain-and loss-of-function results collected from DR^Sert^→OFC and DR^Sert^→CeA neuronal manipulation experiments.

Remarkably, activation of DR^Sert^→OFC neurons significantly enhanced escape behavior (decreased immobility) after CNO application on the day-2 test (Figure 7A). Moreover, ***Tph2*** deletion from DR^Sert^→OFC neurons increased immobility (Figure 7B). Thus, both gain-and loss-of-function experiments indicate that DR^Sert^→OFC neurons promote escape behavior in the forced-swim test. Taken together, these data suggest that activation of DR^Sert^→OFC (but not DR^Sert^→CeA) neurons promotes active coping in face of challenge, and that serotonin release from DR^Sert^→OFC neurons is necessary for this effect.

## Discussion

In mammals, serotonin neurons in the brainstem, in particular the dorsal raphe (DR), project throughout the brain and modulate diverse functions. Several ***a priori*** models could describe the anatomical and functional organization of such a system: (1) All DR serotonin neurons are functionally similar and target downstream regions effectively randomly; collectively they tile the entire target regions to create general serotonin levels, subsequently interpreted by combinatorial expression of serotonin receptors. (2) There may be functionally defined subpopulations, perhaps due to their different inputs, but each subpopulation tile the entire target regions to achieve a similar outcome as Model 1 above. (3) There may be anatomically defined subpopulations, perhaps as a developmental strategy to ensure that all targets are covered, but they are functionally equivalent. (4) The DR is composed of both functionally and anatomically defined subpopulations, and function segregates with anatomical connectivity. While the DR could in principle be simultaneously composed of sub-systems organized according to each of these models, here we provide compelling evidence for the existence of populations that fit Model 4: parallel sub-systems that differ in input and output connectivity, physiological response properties, and behavioral functions. Specifically, we found that CeA-projecting DR serotonin neurons promote anxiety-like behavior, whereas OFC-projecting ones promote active coping in face of challenge, and that these subpopulations have opposing responses to aversive stimuli.

### Anatomical Organization of the DR Serotonin System

How do DR serotoninergic fibers collectively cover their vast target fields? This question can be divided into two related questions: Is there topographic organization of serotonin neurons within the DR with respect to their target fields? For serotonin neurons that project to a specific target, where else in the brain do they also project (i.e., what is their collateralization pattern)? These questions have been addressed with retrograde tracing from target fields (e.g., Jacobs et al., 1978; Fernandez et al., 2016; reviewed in Waselus et al., 2011), or anterograde tracing by limiting the anterograde tracers to subregions of the DR (e.g., Vertes and Kocsis, 1994; Muzerelle et al., 2016) or from genetically defined DR cell types (Bang et al., 2012; Jensen et al., 2008). These pioneering studies led to the proposal that the DR is organized both along the anterior-posterior (Abrams et al., 2004; Commons, 2015) and dorsal-ventral axes (Lowry et al., 2005; Muzerelle et al., 2016) with respect to their target fields, and that individual DR neurons can send collaterals to two or more separate brain regions (Gagnon and Parent, 2014; Waselus et al., 2011). However, these different studies do not fully agree on the details of the topography, perhaps due to several technical limitations. For retrograde labeling, each study usually focused on a small subset of targets, and it is often difficult to quantitatively compare different studies without a common reference brain. For anterograde tracing, the resolution is limited by the spread of injected dyes or the genetic access to subtypes of DR serotonin neurons. Finally, our understanding of the collateralization patterns of DR serotonin neurons is very limited because most of the data were collected by injecting two retrograde tracers into two pre-specified brain regions and observing double-labeled DR serotonin neurons; this method cannot sample the extent of collateralization and often underestimates the true degree of overlap due to incomplete retrograde labeling from each injection site.

Our retrograde tracing from eight diverse target sites, combined with image registration to compare the data in the same reference brain, provides a more comprehensive view of the spatial organization of DR serotonergic projections in the mouse. We did not find a prominent anterior-posterior topographic organization except that entorhinal cortex-projecting serotonin neurons tended to be localized to the posterior DR. We did observe a prominent division along the dorsal-ventral axis in the anterior DR, where ventral and dorsal serotonin neurons preferentially innervate cortical and subcortical targets, respectively, confirming and extending previous reports (Lowry et al., 2005; Muzerelle et al., 2016; Prouty et al., 2017). We further uncovered a strong correlation between cortical projections and Vglut3 co-expression in DR serotonin neurons. An interesting speculation from this finding is that glutamate co-release in cortex endows the DR serotonin system to also regulate cortical circuits for computation in a more rapid time scale using ionotropic glutamate receptors, whereas release of serotonin alone in subcortical circuits primarily serves a slower modulatory function using metabotropic receptors. Indeed, the only ionotrophic serotonin receptors, the HTR3s, are preferentially expressed in cortical and hippocampal interneurons (Barnes and Sharp, 1999). However, it is important to note that our quantitative analyses also revealed that none of these preferences is absolute (Figure 1), highlighting the complexity of the DR serotonin system.

Using our recently developed viral-genetic approaches (Beier et al., 2015; Schwarz et al., 2015) in combination with iDISCO-based tissue clearing and whole-mount imaging (Renier et al., 2016), we were able to describe for the first time whole-brain collateralization patterns of projection-defined DR serotonin neurons. Our detailed analyses of DR^Sert^→OFC and DR^Sert^→CeA axons (Figure 2; Figure S2; Movies S2, S3) demonstrate that they innervate largely complementary targets, consistent with our retrograde studies (Figure 1). These analyses further indicate that the collateralization of individual DR serotonin neurons can be extremely broad— for example, DR^Sert^→OFC neurons also heavily innervate olfactory bulb anteriorally and entorhinal cortex posteriorally, yet highly specific—for example, DR^Sert^→OFC neurons heavily innervate cortical amygdala but avoid nearby basolateral amygdala. These collateralization maps provide a global blueprint of which brain targets are likely coordinately modulated by serotonin, and which targets can be differentially modulated.

cTRIO analyses further reveal that DR^Sert^→OFC and DR^Sert^→CeA neurons receive biased input from specific brain regions (Figure 3). Thus, the input-output architecture of the DR serotonin system differs from that of the locus coeruleus norepinephrine system, where populations of norepinephrine neurons that project to a specific target region also innervate all other regions examined, and receive similar inputs as norepinephrine neurons projecting to any other targets neurons examined (Schwarz et al., 2015). The input-output architecture of the DR serotonin system resembles that of the midbrain dopamine systems (Beier et al., 2015; Lerner et al., 2015), with biased input and segregated output. At least for the DR^Sert^→OFC and DR^Sert^→CeA neurons, the input bias is stronger and output segregation more complete than those of the midbrain dopamine systems, despite the much more extensive collaterization of DR serotonin axons. Future systematic analyses utilizing the methods employed here, ideally supplemented with high-resolution tracing of the axonal arborizations of individual serotonin neurons, will provide a more complete understanding of how the ~9,000 DR serotonin neurons differentially innervate target fields to modulate diverse physiological functions.

### Functional Dissection of the DR^Sert^→OFC and DR^Sert^→-CeA sub-systems

We took two approaches to functionally dissect projection-specific DR serotonin neurons: chemogenetic activation as a gain-of-function approach, and Tph2 (and, by inference, serotonin) depletion as a loss-of-function approach. Because gain-of-function experiments alone may not reflect physiological functions of the system under manipulations, we consider our conclusions stronger if loss-and gain-of-function experiments give opposite results. There are advantages to using Tph2 depletion instead of chemogenetic silencing as a loss-of-function approach. First, chemogenetic silencing requires a higher CNO concentration than does chemogenetic activation; as the active component of CNO may be clozapine (Gomez et al., 2017), which engages several serotonin receptors at high concentrations (Meltzer, 1994), this strategy may ectopically affect serotonin-related behavior. Second, since a sizable fraction of serotonin neurons likely co-release glutamate, Tph2 depletion specifically addresses the function of serotonin in these neurons. A caveat of this approach is that this manipulation is irreversible, and compensatory changes may occur to the circuit during the 2+ weeks between ***AAV_retro_-Cre*** injection and behavioral testing, thus this strategy may underestimate the function of serotonin release. Below, we discuss the functions of DR^Sert^→OFC and DR^Sert^→CeA neurons from these loss-and gain-of-function experiments in conjunction with our input/output mapping and physiological recordings.

#### Locomotion

Our gain-of-function experiments indicate that activation of both DR^Sert^→OFC and DR^Sert^→CeA neurons suppresses locomotion (Figure 5), consistent with previous reports that the DR serotonin system negatively regulates locomotion (Correia et al., 2017; Teissier et al., 2015; Whitney et al., 2016). However, conditional depletion of Tph2 in DR^Sert^→CeA neurons, but not DR^Sert^→OFC neurons, promotes locomotion (Figure 5). These data suggest that the inhibitory effect of DR serotonin neurons on locomotion is at least in part mediated by serotonin release from DR^Sert^→CeA neurons. It is possible that glutamate released from DR^Sert^→OFC neurons also contributes to this effect.

#### Anxiety-like behavior

Both gain-and loss-of-function experiments indicate that DR^Sert^→CeA neurons promote anxiety-like behavior (Figure 6 and Figure S5). DR^Sert^→CeA neurons collateralize to other amygdala nuclei, BNST, and PVH (Figure 2), all of which have been implicated as anxiety-related regions (Calhoon and Tye, 2015). To date, only the function of serotonin innervation in BNST has been extensively investigated in the context of modulating anxiety-like behavior. Fos immunoreactivity is significantly elevated in DR^Sert^→BNST neurons after foot-shock, and optogenetic stimulation of serotonergic terminals in BNST promotes anxiety-like behavior and fear learning (Marcinkiewcz et al., 2016). Our fiber photometry recording in freely moving mice revealed that DR^Sert^→CeA neurons were activated by foot-shocks (Figure 4). Thus, our physiological and behavioral data are consistent with previous findings regarding BNST, and our collateralization studies further indicate that the DR^Sert^→CeA sub-system engages many more anxiety-associated target brain regions. Moreover, cTRIO analysis revealed that the CeA and BNST provided particularly strong input to DR^Sert^→CeA neurons compared to DR^Sert^→OFC neurons (Figure 3). Thus, our data demonstrate that DR^Sert^→CeA neurons promote anxiety-like behavior, likely involving reciprocal connections between the DR and CeA/BNST.

The role of cortical serotonin release in anxiety-like behavior has been extensively studied with pharmacological manipulation and genetic manipulations of serotonin receptors (reviewed in Albert et al., 2014). For example, serotonin receptor knockouts and rescue experiments indicate that cortical HTR2A receptors promote (Weisstaub et al., 2006), whereas cortical HTR1A receptors inhibit (Gross et al., 2002), anxiety-related behavior. However, given the opposing effects of HTR2A and HTR1A in target neurons (enhancing or suppressing excitability), and given the complex distribution of serotonin receptors in different types of excitatory and inhibitory cortical neurons, it is difficult to predict the effect of cortical serotonin release on anxiety based on these results. We found that while chemogenetic activation of DR→OFC neurons did not significantly affect anxiety-like behavior, conditional Tph2 depletion in these neurons enhanced anxiety-like behavior (Figure 6). Since the majority of DR^Sert^→OFC neurons co-express Vglut3 (Figure 1), one interpretation is that activation of DR^Sert^→OFC neurons results in release of both glutamate and serotonin, which could have opposing effects. When serotonin is selectively removed from the pathway, anxiety-like behaviors are promoted. If this were the case, then cortical serotonin release suppresses anxiety.

#### Forced-swim test

Immobility in forced-swim test represents a passive coping strategy in face of challenge, which is often used as an indicator of a depression-like state (Petit-Demouliere et al., 2005). It has recently been reported that chemogenetic activation of DR serotonin neurons reduces immobility in forced-swim test (Teissier et al., 2015). Both our gain-and loss-of-function manipulations in the forced-swim test suggest that DR^Sert^→OFC neurons, but not DR^Sert^→CeA neurons, mediate this effect (Figure 7). Our finding is consistent with a previous study reporting that optogenetic activation of prefrontal cortical neuron terminals in the DR promotes active coping (Warden et al., 2012), given that frontal cortex inputs to the DR preferentially synapse onto serotonin rather than GABA neurons (Weissbourd et al., 2014). Interestingly, DR serotonin neurons directly postsynaptic to frontal cortical inputs localize in the ventral DR (Weissbourd et al., 2014), the origins of DR^Sert^→OFC neurons (Figure 1). Thus, DR^Sert^→OFC neurons and frontal cortex→DR^Sert^ neurons may constitute a reciprocal feedback loop to promote active coping in face of challenge. In addition, our cTRIO data identified biased input to DR^Sert^→OFC neurons from lateral hypothalamus and lateral habenula (Figure 3), both of which are implicated in affective control (Hikosaka, 2010; Stuber and Wise, 2016). It remains a future challenge to determine how DR^Sert^→OFC neurons, and DR serotonin neurons in general, integrate inputs from diverse brain regions to modulate their downstream targets.

In conclusion, our behavioral analyses demonstrate that anatomically segregated DR subsystems have distinct, and sometimes even opposing, functions (Figure 7E). Thus, the DR serotonin system should no longer be viewed or studied as a monolithic population. We have provided technical and conceptual means to further dissect the complexity of the DR serotonin system; such an endeavor will no doubt advance our understanding of neuromodulation and aid in developing effective therapies for psychiatric disorders such as anxiety and depression.

## Acknowledgement

We thank Eliza Adams and Marc Tessier-Lavigne for advice on iDISCO+, Tom Davidson and Karl Deisseroth for advice on fiber photometry, Elizabeth Steinberg and Rob Malenka for advice on behavior, Qi Wu for ***Tph2**^flox/flox^* mice, Alla Karpova for the ***AAV_retro_*** vector, Stanford, Salk Institute, and UNC Viral Core for producing custom viruses, Allen Institute for Brain Science for the reference atlas and Terri Gilbert for the advice on its use, Michael Chen for code files for data analysis, Kevin Beier for DREADD AAVs, Steven Bell for advice on 2D registration, and Rob Malenka, Adi Mizrahi, Andrew Shuster, Laura DeNardo, Jan Lui, Mark Wagner, and Will Allen for critiques on the manuscript. This work was supported by a Hughes Collaborative Innovation Award and a BRAIN initiative grant. L.L. is an HHMI investigator.

## Author Contributions

J.R. designed and performed most of the experiments. D.F. performed iDISCO+ imaging and analysis. J.X. and M.A.H. developed image registration algorithm. C.D.L., K.E.D., C.R., A.P., and Y.S. assisted in experiments or data analyses. B.W. initiated serotonin studies and characterized some of the reagents. R.L.N. produced HSV reagents. L.L. supervised the project. J.R. and L.L. wrote the manuscript with contributions from all coauthors.

---

**Figure S1.**
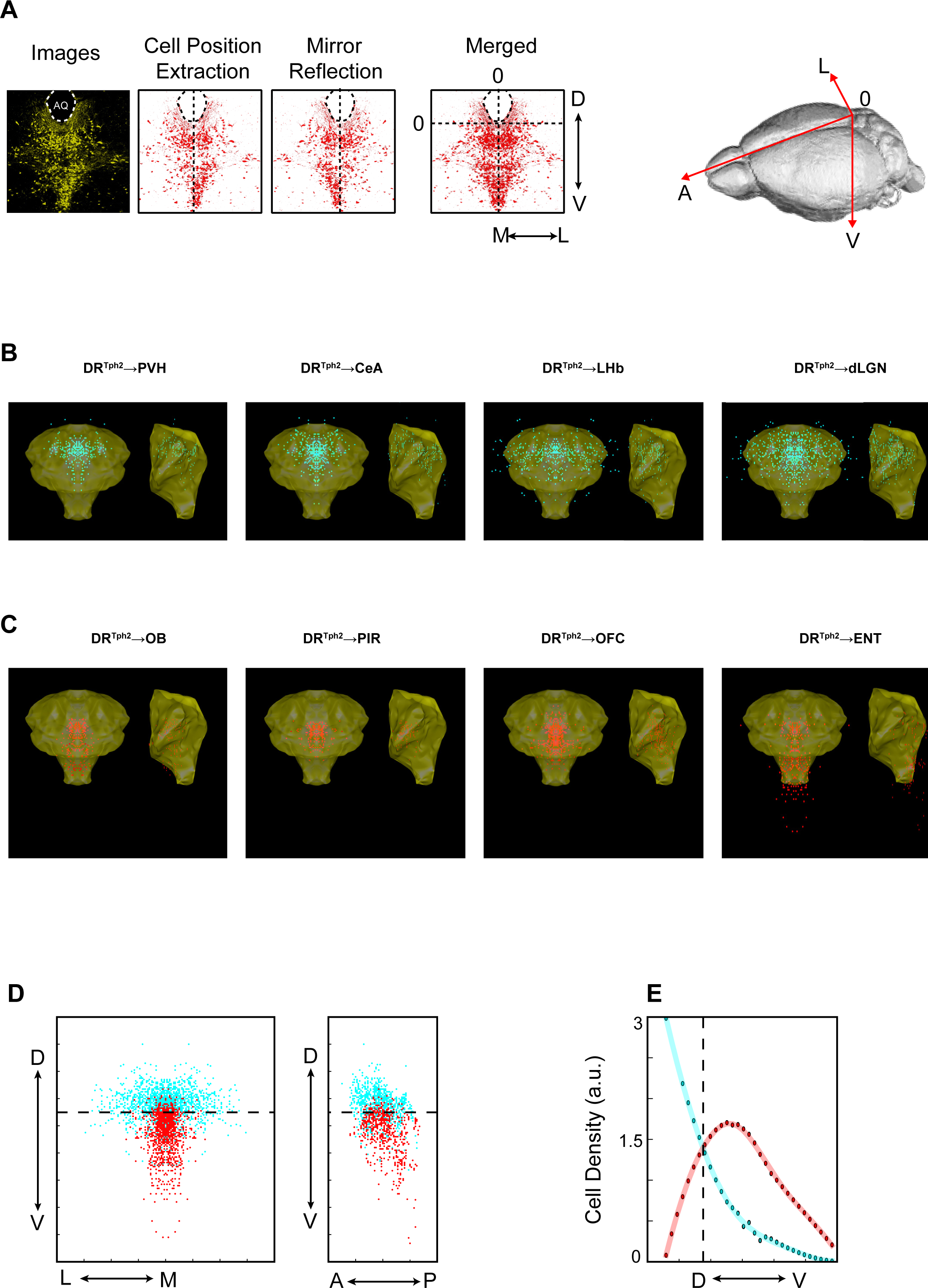
Spatial Organization of DR Serotonin Neurons with Respect to Axonal Projections, Related to Figure 1. (A) Schematic of data collection for 3D reconstruction of DR serotonin neurons. After Tph2 staining, the positions for all the positive neurons were recorded. The line connecting the highest and lowest points of the aqueduct (AQ; dashed ovals) was defined as the midline, and a mirror image was created for each cell across the midline (reflecting the bilateral symmetry of DR serotonin neurons), and then the two mirror images were merged (left four panels). The “zero” along the dorsal-ventral axis refers to the lowest point of AQ for individual slices during data collection. In registered brains, “zero” refers to the brain surface (right panel). (B) Coronal (left) and sagittal (right) view of individual cells’ location (cyan) from four subcortical projecting DR serotonin neuron groups. Yellow represent the surface of the cluster that include all Tph2^+^ cells. (C) Coronal (left) and sagittal (right) view of individual cells’ location (red) from OB and three cortical projecting DR serotonin neuron groups. (D) Coronal (left) and sagittal (right) view of individual cells’ location from a combination of PVH-, CeA-, LHb-, and dLGN-projecting DR serotonin groups (cyan) and a combination of OB-, PIR-, OFC-and ENT-projecting DR serotonin groups (red). Dashed line, 3752 μm below the brain surface. (E) Quantification of cell density along the dorsal-ventral axis. Dashed line shows where the two clusters share the same line density at 3752 μm ventral to the brain surface. The probability for cyan and red cells being dorsal to the 3752-μm deep plane is 75.08% and 15.67%, respectively.

**Figure S2.**
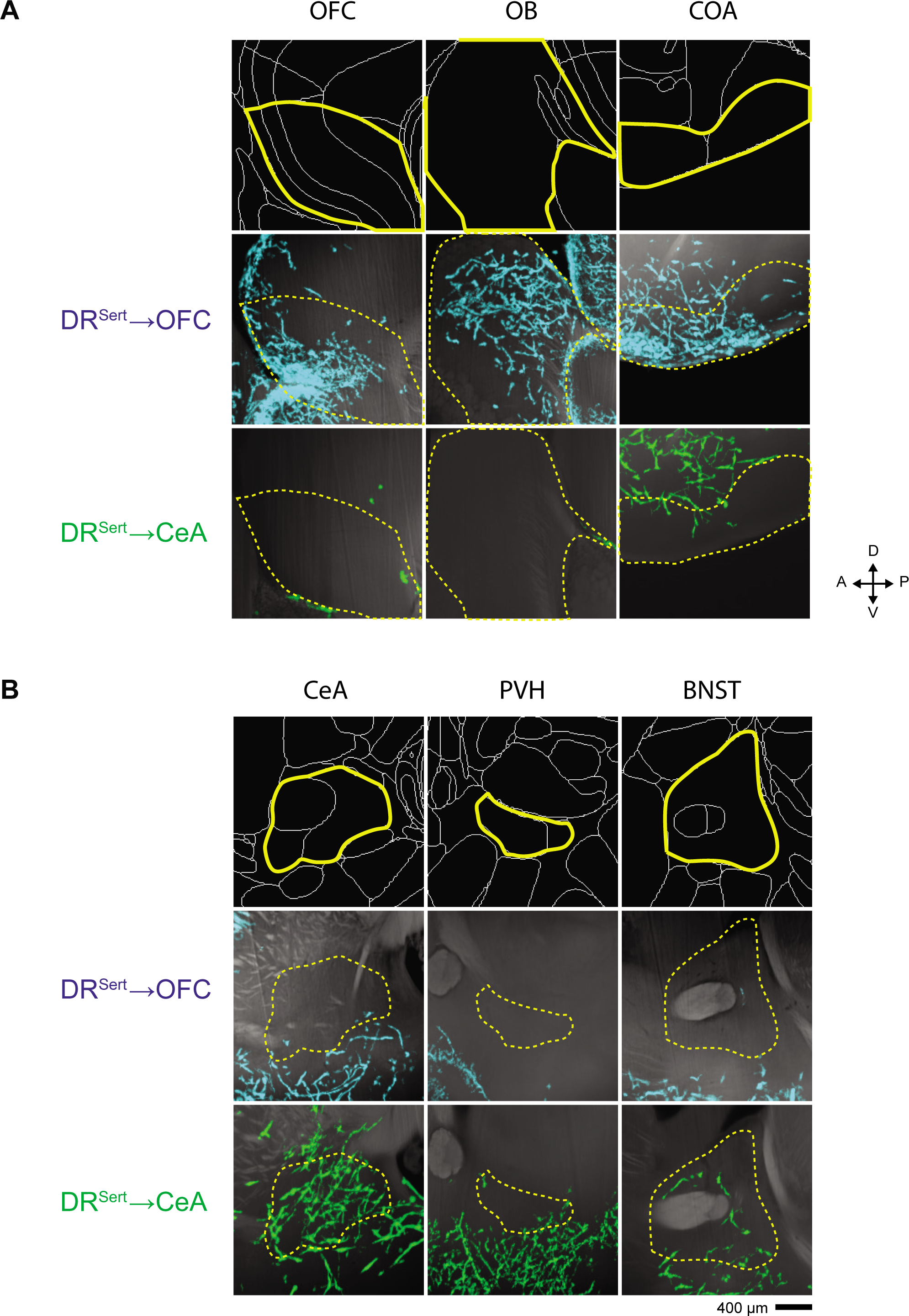
Select Target Regions Highlight the Complementary Nature of Axonal Projections from DR^Sert^→CeA and DR^Sert^→OFC Neurons, Related to Figure 2. (A) DR^Sert^→OFC axons heavily innervate OFC, OB, and the cortical amygdala (CoA) (within the yellow outlines). DR^Sert^→CeA axons largely avoid these regions. (B) DR^Sert^→CeA axons target the CeA as expected, but are also found in PVH and BNST while DR^Sert^→OFC axons are largely absent from these regions. Images of axons are aligned to each brain’s own autofluorescence and both channels are matched to the Allen Common Coordinate Framework. All sections are sagittal with 20 μm in z; scale, 400 mm.

**Figure S3.**
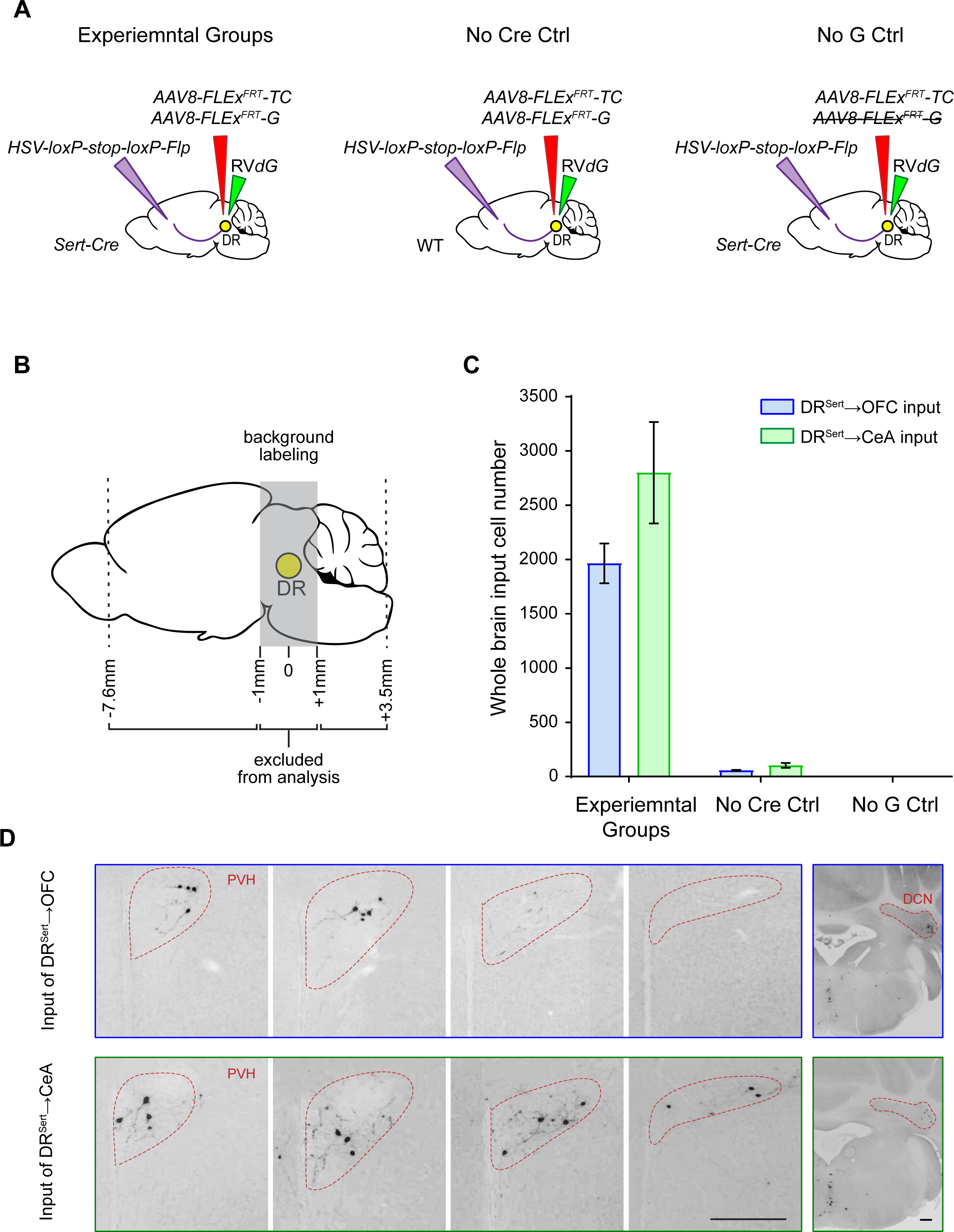
Control Experiments and Additional Images for cTRIO, Related to Figure 3. (A) Schematic of control groups for cTRIO experiments applied on DR. Wild type (WT) mice were used instead of ***Sert-Cre*** mice in No Cre Ctrl, and ***AAV8-FLExFRT-G*** was omitted in No G Ctrl. (B) Schematic of brain regions quantified for input neurons. Regions approximately 1 mm anterior and posterior to the center of the DR were excluded from analysis due to local background labeling from rabies virus (see Weissbourd et al., 2014), including median raphe (MR), ventral tegmental area (VTA), substantia nigra pars compacta (SNc), interpeduncular nucleus (IPN), periaqueductal gray (PAG), locus coeruleus (LC) and several medulla nuclei. (C) Quantification of long-distance background infection in control groups. Y-axis presents total input cell numbers, and X-axis lists group names. (n=9, 8, 4, 4, 4, 4). Error bars, SEM (D) Representatives of GFP labeled ipsilateral input cells in PVH (at four coronal section planes from anterior to posterior; left panels) and in medulla nuclei and DCN (right panels) to OFC- and CeA-projecting DR serotonin neurons. Scale, 250 μm.

**Figure S4.**
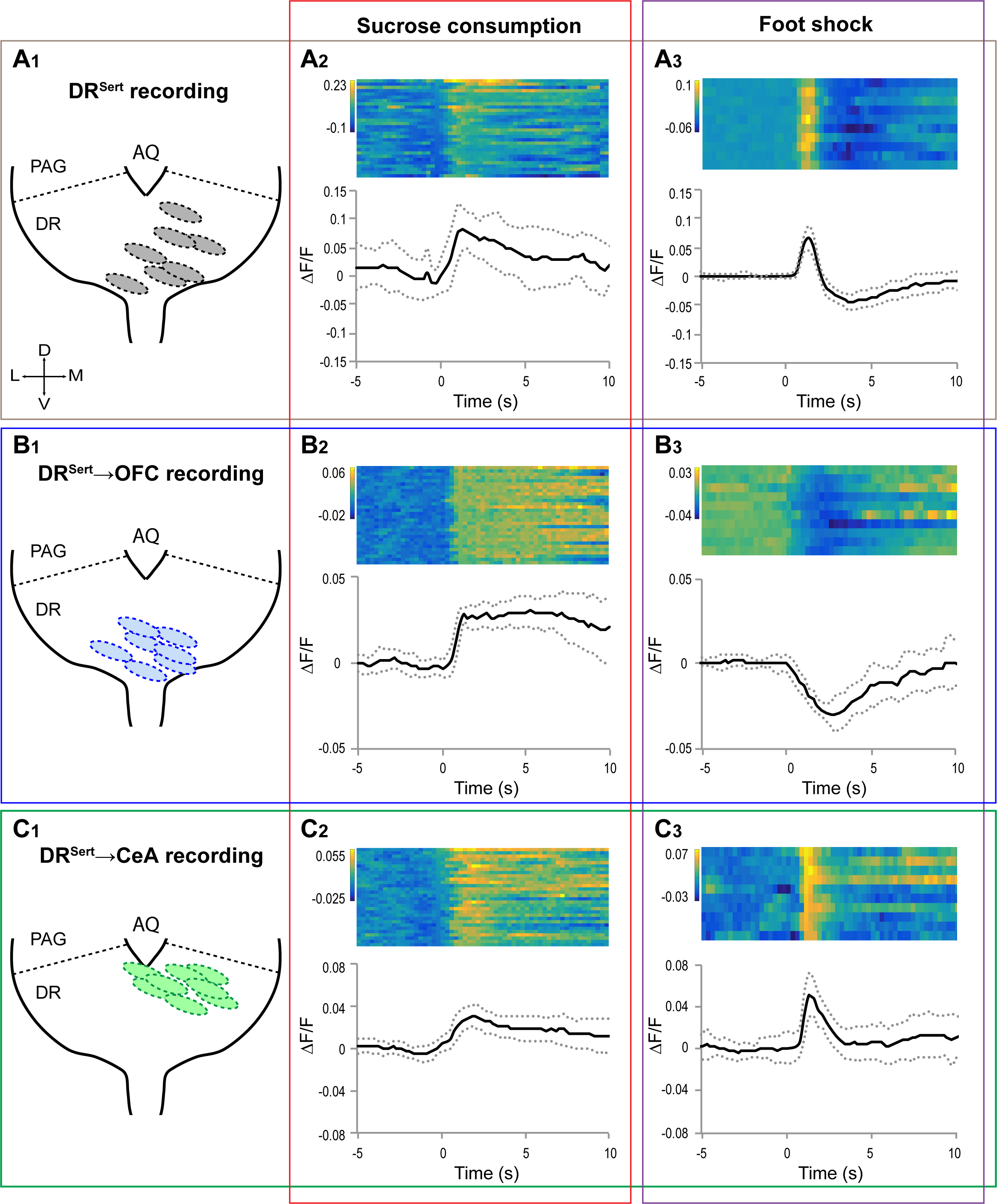
Summary of Optical Fiber Placement and Representative Traces for Fiber Photometry Recording, Related to Figure 4. Fiber photometry recording was performed on DR^Sert^ (A), DR^Sert^→OFC (B), and DR^Sert^→CeA neurons. (A_1_, B_1_, C_1_) Schematic DR diagram showing the end of optic fiber placement (oval) for each mice. AQ, aqueduct. Note that we adjust the fiber implant positions to maximize DR^Sert^→OFC and DR^Sert^→CeA neurons based on data from Figure 1. (A**2**, B**2**, C**2**) Example responses to sucrose consumption observed in one mouse each (the red traces in Figure 4) from the DR^Sert^ (A_2_), DR^Sert^→OFC (B_2_), or DR^Sert^→CeA (C_2_) group. Time 0 is aligned to the initiation of sucrose water licking. Each row of the heat map shows an individual trial. (A3, B3, C3) Example of responses to foot-shock observed in one mouse each (the red traces in Figure 4) from the DR^Sert^ (A_3_), DR^Sert^→OFC (B_3_), or DR^Sert^→CeA (C_3_) group. Time 0 is aligned to the onset of foot-shock. Each row of the heat map shows an individual trial. Solid and dotted lines represent mean ± SEM. Unit for the heap map is AF/F.

**Figure S5.**
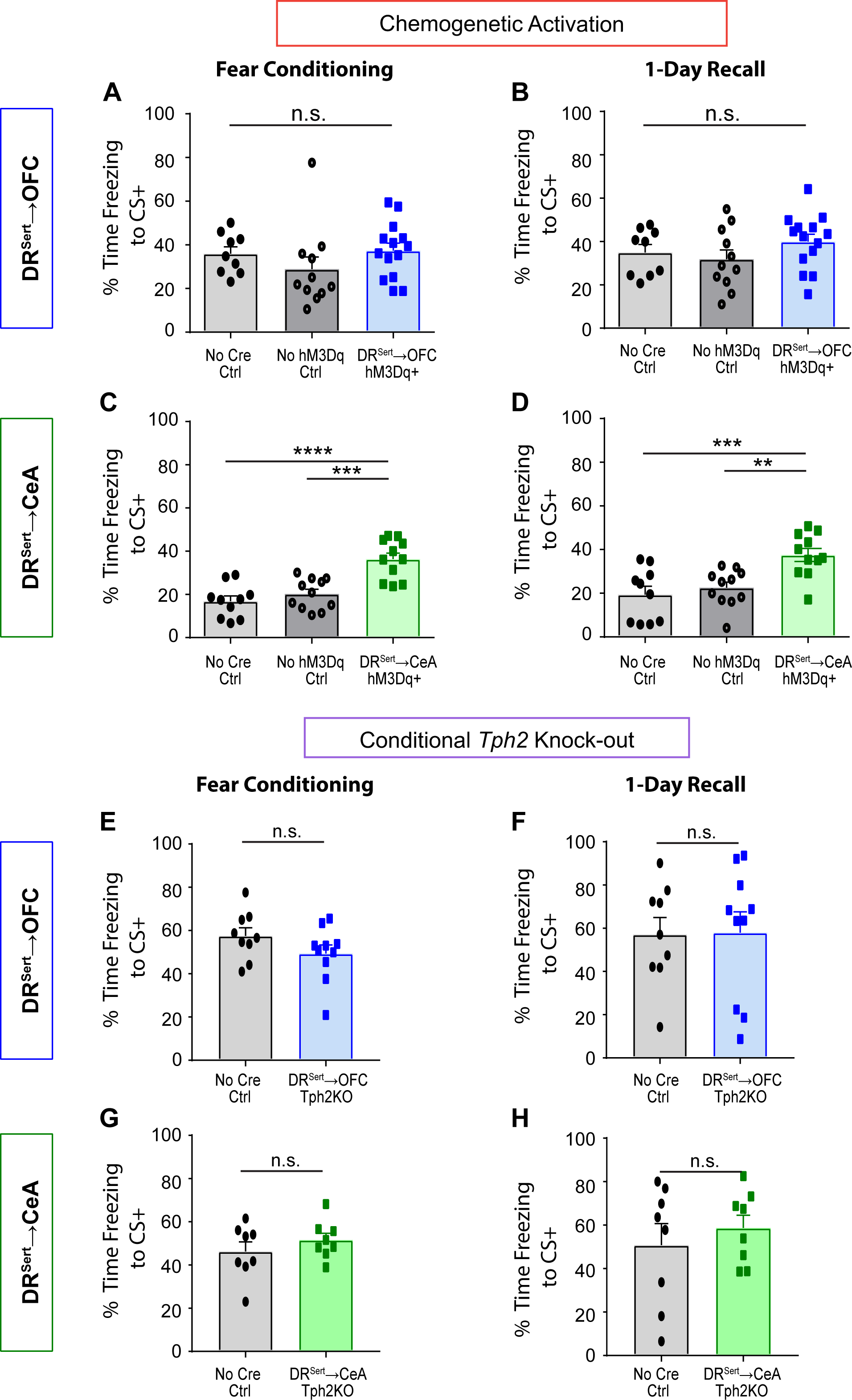
Functional Analysis of CeA- and OFC-projecting DR Serotonin Neurons in Fear Learning and Memory, Related to Figure 6. (A) Chemogenetic activation of hM3Dq+ DR^Sert^→OFC neurons does not affect freezing time to conditioned tone (CS+) during fear conditioning (one-way ANOVA, n= 9, 11, 14). (B) Chemogenetic activation of hM3Dq+ DR^Sert^→OFC neurons does not affect freezing time to CS+ during 1-day recall (one-way ANOVA, n= 9, 11, 14). (C) Chemogenetic activation of hM3Dq+ DR^Sert^→CeA neurons increases freezing time to CS+ during fear conditioning. ***p<0.001, ****p<0.0001 (one-way ANOVA followed by multiple t-tests, Holm-Sidak correction; n= 10, 11, 12). (D) Chemogenetic activation of hM3Dq+ DR^Sert^→CeA neurons increases freezing time to CS+ during 1-day recall. **p<0.01, ***p<0.001 (one-way ANOVA followed by multiple t-tests, Holm-Sidak correction; n= 10, 11, 12). (E) Conditionally knocking out ***Tph2*** from DR^Sert^→OFC neurons does not affect freezing time to CS+ during fear conditioning (two-tail unpaired t-test; n=9, 10). (F) Conditionally knocking out ***Tph2*** from DR^Sert^→OFC neurons does not affect freezing time to CS+ during 1-day recall (two-tail unpaired t-test; n=9, 10). (G) Conditionally knocking out ***Tph2*** from DR^Sert^→CeA neurons does not affect freezing time to CS+ during fear conditioning (two-tail unpaired t-test; n=9, 10). (H) Conditionally knocking out ***Tph2*** from DR^Sert^→CeA neurons does not affect freezing time to CS+ during 1-day recall (two-tail unpaired t-test; n=9, 10).

**Table S1: Allen Brain Atlas IDs and Their Corresponding Names as Identified by the 2017 Common Coordinate Framework, Related to Figure 2**. Regions were selected prior to analysis such that areas defined by individual layers (e.g., cortical layers I-VI), cell identity, and anatomical cardinal directions are collapsed into their parent region. Individual normalized regional densities for each brain are aligned to the heat maps from Figure 2G.

**Table S2: Quantification of cTRIO: Number of Cells in Region (Proportion of Total), Related to Figure 3**. For each sample, the raw cell count for each brain region is displayed in the left column, the fraction of total input for that brain region is displayed in the right column, and the total number of inputs is displayed at the bottom. Medulla nuclei were first calculated individually and then all collapsed into other medulla nuclei expect for the nucleus of the solitary tract (NST).

**Movie S1: 3D Representation of DR Serotonin Neurons’ Spatial Organization with Respect to Axonal Projections and Vglut3 Co-Expression, Related to Figure 1**.

**Movie S2: 3D Rendering of DR^Sert^→OFC and DR^Sert^→CeA Axon Collateralization Patterns, Related to Figure 2**.

**Movie S3: Fly-through Coronal Density Maps of DR^Sert^→OFC (left) and DR^Sert^→CeA (middle) Projections, and A P-value Map (right), Related to Figure 2**.

## STAR METHODS

### KEY RESOURCES TABLE

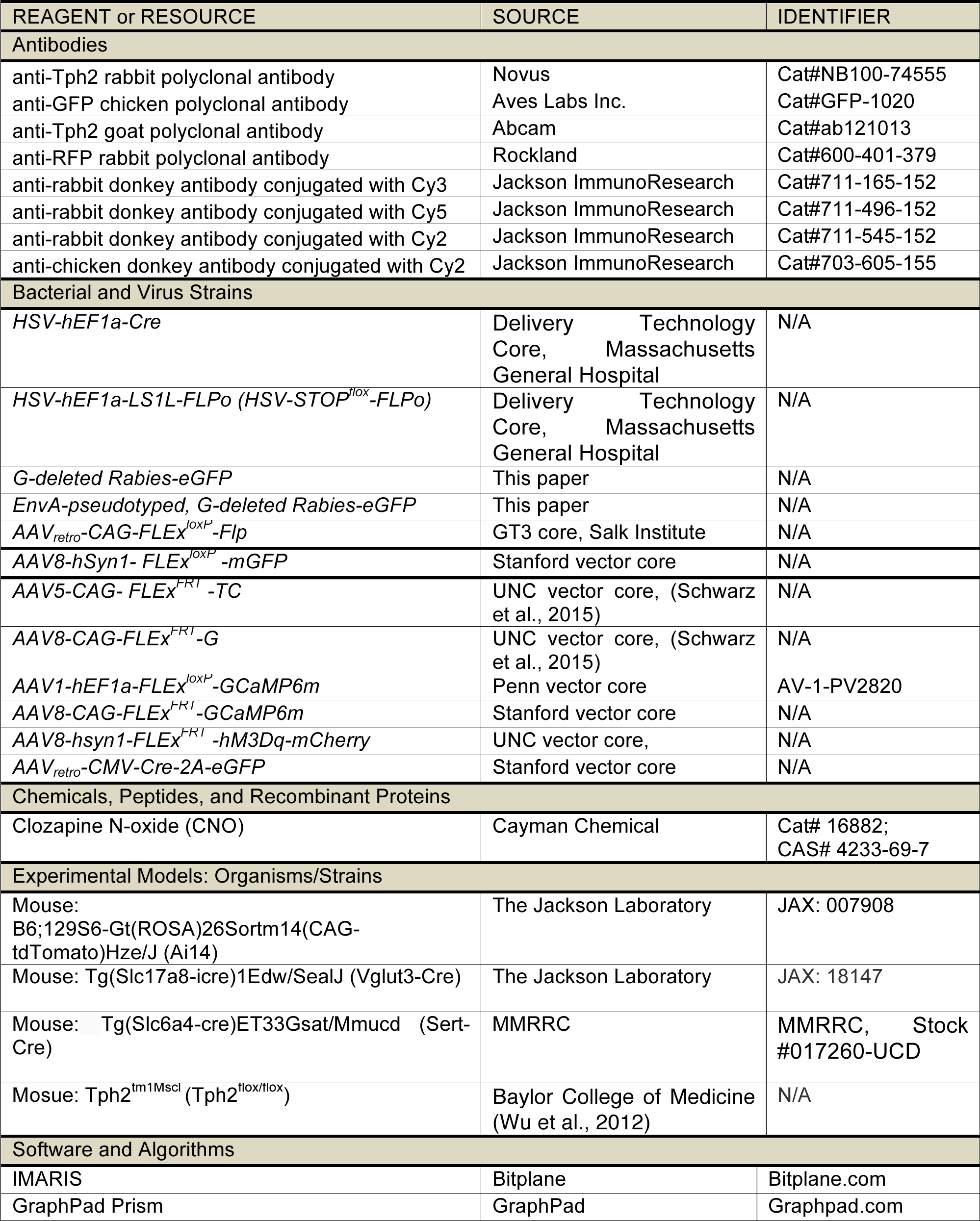
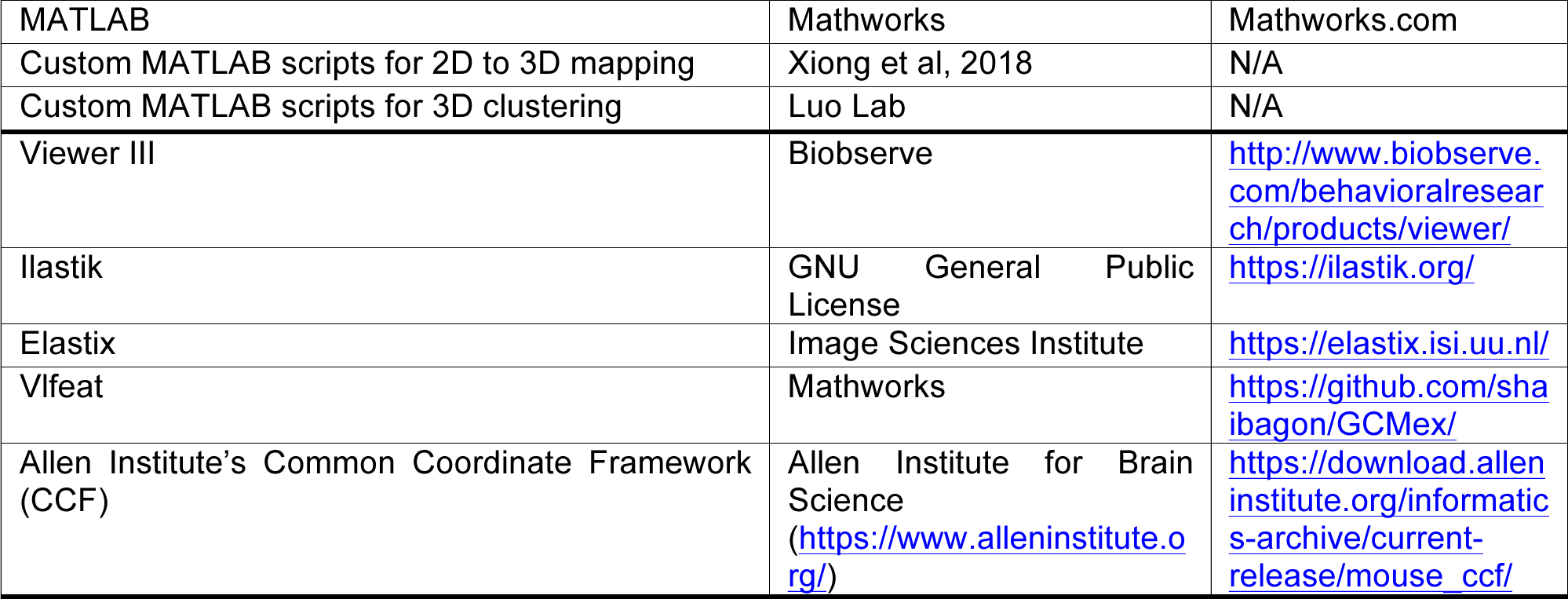

### CONTACT FOR REAGENT AND RESOURCE SHARING

Further information and requests for resources and reagents should be directed to and will be fulfilled by the Lead Contact, Liqun Luo.

## EXPERIMENTAL MODEL AND subJECT DETAILS Animals

### Animals

All procedures followed animal care and biosafety guidelines approved by Stanford University’s Administrative Panel on Laboratory Animal Care and Administrative Panel of Biosafety. For anatomical experiments (Figure. 1-3), male and female mice aged 8-20 weeks on CD1 and C57BL/6J mixed background were used. The ***Ai14*** tdTomato Cre reporter mice (JAX Strain 7914), ***Vglut3-Cre*** (also known as ***Slc18a8-Cre;*** JAX Strain 18147), and ***Sert-Cre*** (MMRRC, Stock #017260-UCD) were used where indicated. For all other experiments (Figure 4-7), male mice aged 8-12 weeks on C57BL/6J background were used when the experiments started. ***Tph2^flox/flox^*** was obtained from Qi Wu (Wu et al., 2012). All mice used in fiber photometry recording were group housed with littermates. All male mice used in gain-and loss-of-function behavioral experiments were individually housed with one female partner. Mice were housed in plastic cages with disposable bedding on a 12 hours light/dark cycle with food and water available ***ad libitum**,* except when placed on water restriction. Experiments were done during the light phase.

## METHOD DETAILS, QUANTIFICATION AND STATISTICAL ANALYSIS

### Stereotaxic Surgeries

Mice were anesthetized with 1.5%-2.0% isoflurane, and placed in a stereotaxic apparatus (Kopf Instruments). For virus injection, the following coordinates (in mm) were used: +4.0 AP, 0.75 ML, −1.5 DV for OB; +2.6 AP, 1.7 ML, −1.7 DV for OFC; + 1.2 AP, 2.8 ML, −4.5 DV for PIR; −3.3 AP, 4.5 ML, −4.5 DV for ENT; 0.2 AP, 0.6 ML, −4.9 DV for PVH; −1.05 AP, 2.86 ML, −4.55 DV for CeA; −1.4 AP, 0.4 ML, −2.6 DV for LHb; −2.3 AP, 2.6 ML, - 2.7 DV for dLGN; −4.3AP, 1.10 ML, −2.85 DV for DR, with 20° ML angle. (AP is relative to bregma; DV is relative to the brain surface when AP is −1.0). For fiber photometry experiments, a fiber optic cannula was implanted over the DR through the same hole as made for the virus injection. To reduce autofluorescence artifacts and maximize light collection, cannulae (special order from Doric Lenses) were fabricated using 0.48 NA 400 p,m BFH48-400 fiber, non-fluorescent epoxy and metal 2.5mm ferrules. Cannulae were fixed to the skull using dental cement (Parkell, C&B metabond). After surgery, mice were allowed to recover until ambulatory on a heated pad, and then returned to their homecage.

### Viruses

Viruses were injected with the following volumes and titers:

***HSV-hEF1a-cre**,* 2 x 10^9^ infectious units/ml;

OB, 500 nl; OFC, 750 nl; PIR, 500nl; ENT, 500 nl; PVH, 300 nl; CeA, 300 nl; LHb, 300 nl; dLGN, 500 nl.

***HSV-hEF1a-LS1L-FLPo**,* 5 x 10^9^ infectious units/ml;

OFC, 750 nl; CeA, 300 nl. ***eGFP-expressing G-deletedRabies Virus**,* 1 x 10^9^ gc/ml;

OB, 500 nl; OFC, 750 nl; PIR, 500 nl; ENT, 500 nl; PVH, 300 nl; CeA, 300 nl; LHb, 300 nl; dLGN, 500 nl.

***EnvA-pseudotyped, eGFP-expressing G-deleted Rabies**,* 1 x 10^9^ gc/ml;

DR, 500 nl.

***AAVretro-CAG-FLEx^loxP^-Flp**,* 6.9 x 10^12^ gc/ml;

OFC, 750 nl; CeA, 300 nl.

***AAV8-hSyn1-FLEx^loxP^-mGFP**,* 2.9 x 10^13^ gc/ml;

DR, 500 nl.

***AAV5-CAG-FLEx^FRT^-TC**,* 2.6 x 10^12^ gc/ml, ***AAV8-CAG-FLEx^FRT^-G**,* 1.3 x 10^12^ gc/ml;

DR, 500 nl;

***AAV1-hEF1a- FLEx^loxP^-GCaMPôm**,* 1 x 10^13^ gc/ml;

DR, 500 nl;

***AAV8-CAG- FLEx^FRT^-GCaMP6m**,* 1.8 x 10^13^ gc/ml;

DR, 500 nl;

***AAV8-hsyn1-FLEx^FRT^-hM3Dq-mCherry**,* 6.2 x 10^12^ gc/ml;

DR, 500 nl;

***AAV_retro_-CMV-Cre-2A-eGFP**,* 8.7 x 10^12^ gc/ml;

OFC, 750 nl; CeA, 300nl.

### Histology and Imaging

Animals were perfused transcardially with phosphate buffered saline (PBS) followed by 4% paraformaldehyde (PFA). Brains were dissected, post-fixed in 4% PFA for 12-24 hours, and placed in 30% sucrose for 24-48 hours. They were then embedded in Optimum Cutting Temperature (OCT, Tissue Tek) and stored at in the −80°C freezer until sectioning. For the antibody staining in Figure 1, 50-μm sections containing DR were collected onto superfrost Plus slides to keep the anterior to posterior sequence. All the working solutions listed below were added with 0.2% NaN_3_ to prevent microbial growth. The slices were then washed 3×10 min in PBS and pretreated with 0.5mM SDS in 37°C incubator overnight. And then they were blocked for 4 hours at room temperature in 10% normal donkey serum (NDS) in PBS with 0.3% Triton- X100 (PBST). Primary antibody (Novus, rabbit anti-Tph2) was diluted 1:1000 in 5% NDS in PBST, and incubated for 24 hours at room temperature. After 3×10 min washes, secondary antibody was applied for 6 hours at room temperature (donkey anti-rabbit, Alexa-647 or Alexa- 488, Jackson ImmunoResearch), followed by 3×10 min washes in PBST. And then slices were stained for NeuroTrace Blue (NTB, Invitrogen). For NTB staining, slides were washed 1×5 min in PBS, 2×10 min in PBST, incubated for 2-3 hours at room temperature in (1:500) NTB in PBST, washed 1×20 min with PBST and 1×5 min with PBS. Sections were additionally stained with DAPI (1:10,000 of 5 mg/mL, Sigma-Aldrich) in PBS for 10-15 min, and washed once more with PBS. The slides were mounted and coversliped with Fluorogel (Electron Microscopy Sciences). These sections were then imaged using a Zeiss 780 confocal microscope, and images were processed using NIH ImageJ software. After that, whole slides were then imaged with a 5× objective using a Leica Ariol slide scanner with the SL200 slide loader.

For long-range tracing analysis in cTRIO experiments (Figure 3), whole brain consecutive 50-μm coronal sections except DR containing ones were collected and NTB stained as described above. For DR containing slices in Figure 3-7, staining was applied to floating sections. Primary antibodies (Novus, rabbit anti-Tph2, 1:1000; Rockland, rabbit anti-RFP, 1:1000; Abcam, goat anti-Tph2, 1:500; Aves Labs Inc., chicken anti-GFP, 1:2000) were applied for 48 hours and secondary antibodies 12 hours at 4°C.

### 2D Registration

For 2D registration (Figure 1 and S1), whole-slide images of scanned slides were imported into custom Matlab software to segment images into individual brain sections based on the NTB stain. To accelerate processing, the full resolution images (xy-resolution =1.29 μm/pixel) were initially down-sampled by a factor of 32 in both x- and y- dimensions. Segmentation included the application of a mask fit to the edge of each section to remove all image features outside the section. Background subtraction and contrast enhancement of the NTB channel were then applied. The processed NTB images for each section were then serially analyzed using a combination of automated and manual methods. To estimate the sectioning angle difference between samples and Allen reference brain (Allen Institute for Brain Science, 2015, http://brain-map.org/api/index.html), we assume parallel cutting angle throughout the sectioning for each brain. To generate standard atlas regarding different angles, the atlas was rotated, re-sectioned into coronal slices, and re-index the slices in order. The histological sequence from each sample was compared with the newly generated re-sectioned atlas slices. Every third slice of the experimental brain was selected automatically to quantitatively evaluate for the cutting angle difference, while severely damaged slices were skipped if necessary. The images from each group were first brought to the same coordinates with a similarity transformation estimated with the Umeyama method based on contour point correspondence generated by Shape Context. Slices were further rescaled in the horizontal and vertical direction to accommodate the global deformation. Features identified by Histogram of Oriented Gradients (HOG) were then extracted from both images, and the L2 norm of HOG difference was used as the similarity metric. The difference between two images is measured as a scalar, which is the summation of the HOG difference over all blocks. Matching slice index difference of half brains was used to determine the cutting angle. Matching slice indices were then interpolated linearly to identify the best matching atlas section for each sample slice in the experimental brain. All the experimental slices were registered nonrigidly to their computed corresponding slice in the optimally rotated atlas to build a pixel-wise mapping from the 2D slice sequence to the reference volume. We augmented the Markov random field (MRF) approach to model brain tissue coherency. We further made improvement based on the data-specific properties of our experimental dataset including segmenting the aqueduct with a convolutional neural network and local warping them with thin plate spline (TPS). For a more detailed description of this procedure see Xiong et al. (2018).

### 3D Reconstruction and Clustering of DR Neurons

To construct the volume presenting DR serotonin neurons, slices containing the Tph2-positive neurons from four animals were registered to Allen’s reference atlas. The line connecting the highest and lowest points of the aqueduct was defined as the midline, with a “zero” value along the medial-lateral axis. To reflect the bilateral symmetry of the DR serotonin system, mirror images were created for each cell across the midline plane (Figure S1A). All the cells’ 2D positions were determined automatically by custom Matlab program employing k-means, following with 3D registration. DBSCAN was performed to cluster the combined data and establish a 3D surface of the DR serotonin system that covered ~97% of Tph2-positive neurons using custom software. Delaunay Triangulation was then performed on the clusters outputted from DBSCAN to define the boundary. Catmull Clark subdivision was then applied to the boundary to finalize the shape of 3D clusters.

For each brain with specific retrograde injections and Tph2/Vglut3 dual labeling neurons, the 2D positions were determined manually and registered to the same reference atlas, allowing cross comparison of the data from the DR of different brains. DBSCAN was performed to cluster individual group and establish each 3D surface that covered ~85% of the neurons. Further processing followed the procedures described above.

### Cell Density and Line Density

Cell density at location D is a function to calculate cell numbers located in a 3D coordinate system defined by ***x*** (axis M→-L), ***y*** (axis D→-V), z (axis A→-P), denoted as ***D(x,y**,*z). Define ***N(x_0_,y_0_,z_0_)*** to be the number of cells located in space defined by ***x < x_0_,y < y_0_, z < z_0_.***

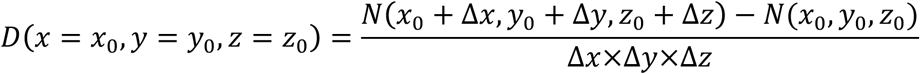

Cell linear density along ***y*** at location ***y_0_*** is defined as

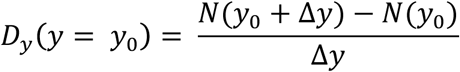

***N(y***_0_) means the number of cells at location defined by **y** < **y_0_**, i.e. ventral to the brain surface at **y_0_** nm.

### iDISCO-based Whole-Brain Axon Tracing

Brains were perfused, dissected, and processed according to the iDISCO+ pipeline as previously described (Renier et al., 2016). Whole brains were processed in 5-ml volumes, labeling in 1:2000 anti-GFP (Aves, GFP-1020) for 10 days and secondary Alexafluor 647 (Jackson Immunoresearch) for 7 days. Images were collected with a LaVision Lightsheet UltramicroscopeII at 0.8× magnification using 640 nm and 488 nm imaging lasers and a z-step size of 3 μm. The working distance of the microscope allowed visualization of the right hemisphere of each brain in the sagittal plane with an approximate 6 mm imaging depth. The image stack of GFP+ axons in the 640-nm channel was first processed with a series of high pass filters to reduce background noise and striping artifacts generated by shadows from the lightsheet. A 2D pixel classifier was trained in Ilastik using 2-5 images from each of 8 brains. Autofluorescent fiber tracts were separated from labeled axons with a second pixel classifier. Contiguous 3D objects were classified in Matlab according to volume, solidity, orientation, intensity, and proximity to remove artifacts with similar properties. The image stack of autofluorescence in the 488 nm channel was aligned to a reference brain generated by serial two photon tomography that was co-registered to the Allen Institute’s Common Coordinate Framework (CCF) (Kim et al., 2015). subsequently, the processed stack of axons was transformed to the same coordinates. Registration and transformation were performed using the Elastix toolbox (Klein et al., 2010; Shamonin et al., 2013). Voxels classified as axons were equally thresholded in all brains and counted by regions as described in the 2017 CCF. Within the Allen’s hierarchy of brain areas, regions distinguished solely by layers or anatomical location were collapsed into their “parent” region (e.g., Layers 1-6 of both dorsal and ventral anterior cingulate area are labeled as “anterior cingulate area”). These decisions were made prior to analysis and were agreed to by four separate anatomical experts. Reported values of axonal labeling density for individual brain regions are normalized both to the volume of the region itself and the total labeling density for that sample to eliminate variability from injection volume. Fiji and Imaris software were used to generate images.

### cTRIO Experiments

Mice were anaesthetized and injected with 500 nl of a 1:1 mixture of ***AAV8 CAG-FLEx^FRT^-G*** and AAV5 ***CAG-FLEx^FRT^-TC*** into the DR, and also injected either with 750 nl ***HSV-STOP^flox^-Flp*** into ipsilateral OFC or 300 nl into ipsilateral CeA using coordinates described above. After recovery, mice were housed in a BSL2 facility. Two weeks later, 500 nl RV***dG*** was injected into the DR using the procedure described above. After recovery, mice were housed in a BSL2 facility for 5 days before euthanasia.

Cell counting was performed manually using Fiji. For quantifications of subregions, boundaries were based on the Allen Institute’s reference atlas (Lein et al., 2007) with consultation of Franklin and Paxinos (2013). The infralimbic cortex and medulla are as defined in the Allen atlas; for medulla, sections anterior to the appearance of the DR were omitted due to possible local background (Figure S3). For counts of thalamic subregions, we were conservative while counting sections that border midbrain nuclei, so our counts may underestimate posterior thalamic subregions. We did not adjust for the possibility of double counting cells from consecutive sections, which likely results in overestimates, with the extent depending on the size of the cells in the regions quantified.

### Fiber Photometry

Fiber photometry was performed using modulated 405 nm and 490 nm LEDs (). The light path was coupled to a 0.53-NA, 400-μm optical fiber patch cord, which was then coupled to the fiber implant in each mouse. Behavioral data from the operant system was synchronized to the fluorescence data using a TTL pulse at the start of each session. Signals were digitized using a digital signal acquisition board, demultiplexed using a software lock-in amplifier, and then low pass filtered to 30 Hz before saving to disk at 381 samples/s. At the start of each recording session, fluorescence values in the 490-nm and 405-nm channels were approximately matched to ensure accurate fitting and subtraction during analysis. Before recording during behavior, mice were screened to check for high GCaMP expression. For the lever-pressing experiments, GCaMP-expressing mice were trained to drink in the operant box after lever-pressing. After reaching proficiency, mice were tested with the following protocol: after 48 hours water-restriction, mice were allowed to lever press to obtain 5% sucrose reinforcements for at least 25 trials. The fluorescence signals, reinforcements, and licks were recorded throughout the session. For the foot-shock experiments, GCaMP-expressing mice were first habituate to the shock box for 15 minutes. Then in a 15-min recording session, 5 tones were delivered, each marked by a TTL pulse. 24 hours later, mice were exposed to 10 0.5-mA and 1-sec electric shocks. The start of each shock was marked by a TTL pulse.

Fiber photometry data were analyzed in MATLAB. Each channel was loaded and resampled to 3.81 Hz. The 405-nm channel was scaled to the 490-nm channel using a least-squares fit, and then ΔF(t)/F0 = (F490(t) - scaledF405(t)) / scaledF405(t) was computed and smoothed with 1.9 s moving average filter. Finally, the median ΔF(t)/F0 from the 5-min baseline period (prior to licking, or stimulus delivery in each experiment) was subtracted from the entire trace.

### Drug Administration

Clozapine N-oxide (CNO; Cayman Chemical, Item No. 12059) was dissolved in 0.4% DMSO and 0.9% NaCl to achieve 1 mg/kg body weight when administered by ***intraperitoneal injection.*** All CNO injections occurred 40 min before the onset of behavioral tests (Teissier et al., 2015).

### Behavior Assays

The behavioral assays described below are in the order of assay performance. Each assay was segregated from the last one for at least 5 days. No animals were excluded from behavioral experiments except the one died before all the assays were completed.

#### Open field

The open field apparatus consisted of a 50 cm x 50 cm clear Plexiglas arena. Mice were acclimated to the experimental test room for at least 30 min prior to testing. To start a session, a mouse was placed into the center of the arena and allowed to freely explore for 10 minutes with video recording. The total distance traveled (m), time spent in center (25 cm x 25 cm) (s) and center entry were analyzed later from recorded video automatically by software (Biobserve).

#### Elevated plus maze

The elevated plus maze apparatus consisted of two open and two closed arms extended out from a central platform. Each arm of the maze was 30-cm long and 5-cm wide. The maze surface was 85-cm above the floor. Each mouse was placed in the same position on the open arm of the maze at the beginning of the assay, facing the center, and allowed to explore the apparatus for 5 minutes. The number of open and closed arm entries as well as the total time spent in open and closed arms were analyzed later from recorded video automatically by software (Biobserve).

#### Auditory fear conditioning

Mice were habituated to the conditioning chamber and tones for 15 min per day for 3 days. On the fourth day, animals for gain-of-function experiments received one CNO injection 40 min before fear conditioning. The fear-conditioning chamber consisted of a square cage (18 x 18 x 30 cm) with a floor wired to a shock generator and a scrambler, surrounded by an acoustic chamber (Coulbourn Instruments). We used two tones in a differential auditory fear conditioning protocol (CS+: 4 kHz, 30 sec, ~75 dB and CS-: 16 kHz, 30 sec, ~75 dB). The protocol consisted of 4 baseline tones (2 CS+, 2 CS-, interleaved), followed by interleaved presentations of 8× CS+ that co-terminated with a 1-sec, 0.25-mA foot-shock for gain-of-function experiments and 0.5-mA for loss-of-function experiments, and 4× CS- that were not paired with a shock. During a 1-day memory retrieval session, animals returned to the conditioning chamber and were presented with interleaved 8× CS+ and 4× CS-.

#### Forced-swim test

Mice were placed for 6 min in a plastic cylinder (height: 25 cm; diameter: 18.5 cm) filled with water (15 ± 1°C) to a depth of 14 cm. The water depth was adjusted so that the animals were forced to swim or float without their hind limbs touching the bottom. The session was videotaped and analyzed by two trained researchers individually afterwards on the computer blind to genotype. Duration of immobility (the time during which the subject made only the small movements necessary to keep their heads above water) was scored by averaging the results from the two researchers. A two-day forced-swim test was applied. For gain-of-function experiments, mice only received one CNO injection 40 min before the 2^nd^ day test.

### Quantification and Statistical Analysis

All statistical tests and data analyses were performed using MATLAB and GraphPad Prism. Full details of each statistical test used are described in each figure legend. Significance was defined as p < 0.05. Sample sizes were chosen based on those used in previous papers.

